# Neuropeptidergic transmission shapes emergent properties of prefrontal cortical circuits underlying learning

**DOI:** 10.1101/2025.05.13.653840

**Authors:** Miguel A. Arenivar, Nathanyal Ross, Rodolfo J. Flores, Sultan Zakariya, Zachary Wang, Hector E. Yarur, Rufina Kore, Wilma V. Richiez-Mateo, Aaron Limoges, Huikun Wang, Hector Bravo-Rivera, Diana Agbor-Enoh, Juan Enriquez-Traba, Xinglong Gu, Bruno B. Averbeck, Yulong Li, Hugo A. Tejeda

## Abstract

The prefrontal cortex (PFC) is essential for top-down control of affect and its dysfunction is implicated in many psychiatric disorders. Inhibitory interneurons expressing somatostatin have been implicated in cognition, affect, and disease. However, somatostatin’s function as a neuropeptide transmitter remains unclear. Here, we investigated the contribution of somatostatin neurotransmission in differentiating between salient (rewarding or aversive) and neutral outcomes. Monitoring somatostatin release and somatostatin receptor antagonism revealed time-dependent regulation of outcome-specific associative learning during acquisition. We found that somatostatin transmission enables configural representations incorporating a salient, aversive outcome in prefrontal cortical neurons, and a threat-driven shift in population-level network activity *in-vivo*. These findings show a novel role for somatostatin, an interneuron “cellular marker”, signaling in shaping learning and emergent network dynamics. Further this framework revealed that reduced somatostatin neuropeptidergic transmission may impair top-down control of affective behaviors observed in mental health disorders, suggesting potential new avenues for therapeutic intervention.

## Introduction

The prefrontal cortex (PFC), is critical for top-down control of adaptive behavior, including selection of appropriate behaviors to neutral and salient outcomes (Averbeck & Murray, 2020; Cummings et al., 2022; Friedman & Robbins, 2022; Miller & Cohen, 2001). Emergent properties of PFC circuits including changes in neuronal representations of task and behavioral features at the population-level underlie executive control of behavior (Averbeck et al., 2006; Fusi et al., 2016; Panzeri et al., 2022; Tye et al., 2024). The PFC is composed of microcircuits containing excitatory pyramidal cells and inhibitory interneurons, which are classified into distinct sub-populations based on morphological properties, intrinsic physiology, calcium-binding proteins, and neuropeptide expression, including somatostatin (SST), neuropeptide Y-, parvalbumin-, and vasoactive intestinal peptide-positive interneurons (Anastasiades & Carter, 2021; Kepecs & Fishell, 2014; Letzkus et al., 2015; Wu et al., 2023). Though connectivity of distinct interneurons in PFC circuits through fast inhibitory GABAergic synapses has been studied, a critical gap remains in understanding the physiological role of neuropeptides.

Neuropeptides, such as SST, act as inter-cellular signaling molecules via their actions on G-protein coupled receptors (GPCRs), which are expressed in both the human and rodent cerebral cortex (Casello et al., 2022; Fee et al., 2017; Song et al., 2021). For example, despite serving as a canonical “marker” of SST interneurons, the consequences of SST neuropeptidergic transmission in the mPFC are not well understood. Limited physiological studies have demonstrated that SST can have complex cellular actions on neocortical neurons. This includes direct and indirect hyperpolarization or depolarization of pyramidal and putative non-pyramidal neurons in the PFC (Brockway et al., 2023), and suppression of excitatory inputs innervating fast-spiking parvalbumin cells in the primary visual cortex (Song et al., 2020). Thus, SST neuropeptidergic transmission may shape neuronal representations in cortical networks *in-vivo* in a complex manner, rather than causing widespread increases or decreases in cortical activity. Intraventricular infusions of exogenous SST peptide or SST-receptor (SSTR) 2/3 agonists (Engin et al., 2008; Engin & Treit, 2009) or intra-mPFC SSTR2/3 agonist (Brockway et al., 2023) produced antidepressant-like or anxiolytic effects, respectively. Microinjection of SST or a SSTR receptor antagonist in the primary visual cortex improved and impaired visual perception, respectively (Song et al., 2020). Interestingly, dysregulation of PFC SST expression is implicated in numerous psychiatric disorders, including depression, bipolar disorder, and schizophrenia (Beneyto et al., 2012; Fung et al., 2014; Hashimoto et al., 2008; Lin & Sibille, 2015; Morris et al., 2008; Pantazopoulos et al., 2017; Seney et al., 2015; Tripp et al., 2011). However, studies identifying the causal role of endogenous SST release in regulating top-down control of behaviors relevant to mental health disorders and neuronal representations are limited in large part due to the lack of tools to monitor and manipulate SST release and its impact on cortical circuits *in-vivo*.

In this study, we used a combination of genetic and viral approaches, *in-vivo* fiber-photometric recordings using a novel genetically-encoded SSTR-based fluorescent sensor, single-cell Ca^2+^ imaging, and behavior to determine the role of endogenous SST transmission in the medial PFC (mPFC) in discrimination of salient and neutral outcomes and their predictors. We demonstrate that mPFC SST (mPFC^SST^) neuropeptidergic transmission is critical for the discrimination of both threats and rewards with neutral outcomes. We uncover SST neuropeptide release dynamics and demonstrate a time-dependent role of SST neuropeptide transmission during discriminative threat learning. Overall, we show that SST-peptidergic transmission is necessary for mPFC circuits to generate configural representations in mPFC networks associated with discrimination of neutral and salient outcomes.

## Results

### mPFC^SST^ knockdown impairs cued auditory threat discrimination

We were interested in the potential contribution of SST in the mPFC in discrimination and developed a method to selectively eliminate SST (mPFC^SST^-cKO). We injected SST-loxP mice with intra-mPFC AAV-Cre-GFP (Huang et al., 2018; Fig. 1A). As expected, SST peptide immunoreactivity was decreased in mPFC^SST^-cKO mice relative to control (WT) mice injected with AAV-Cre-GFP (Fig. 1A). We next used cued auditory threat discrimination to examine how mPFC^SST^-cKO impacts discrimination between a cue predicting a negatively valenced outcome and a neutral cue associated with no outcome (Fig. 1B). During conditioning, mice were exposed to an auditory cue co-terminating with a footshock (CS^+^) and a neutral tone without consequence (CS^−^). Freezing, a passive defensive behavior elicited by threats, was similar in control and mPFC^SST^-cKO mice during baseline habituation and threat acquisition (Fig. S1A-D). As expected, during cued-threat recall where the CS^+^ and CS^−^ were presented in a different context not associated with footshock, control mice exhibited robust freezing to the CS^+^, but not the CS^−^ (Fig. 1 C, D). In marked contrast, mPFC^SST^-cKO mice displayed robust freezing to both the CS^+^ and CS^−^ (Fig. 1 C, D). This suggests that mPFC^SST^ signaling does not modulate retrieval of threat-associated memories driven by the CS^+^, but instead promotes discrimination between cues associated with threats and neutral outcomes. Importantly, enhanced freezing was not observed during the pre-CS or inter-trial period during threat recall test day (Fig. 1C, D; Fig. S1E), demonstrating that enhanced freezing to the CS^−^ in mPFC^SST^-cKO mice is not a consequence of generalized freezing after threat conditioning procedures. Of note, locomotor activity elicited by footshock on both conditioning days was not different between control and experimental mice (Fig. S1F,G). To control for potential effects of the SST-loxP strain, we injected separate groups of WT or SST-loxP mice with either unilateral or bilateral AAV-Cre-GFP (Fig. 1E). In contrast to bilateral mPFC^SST^-cKO, unilateral mPFC^SST^-cKO displayed robust and minimal freezing to the CS^+^ and CS^−^, respectively, similar to WT controls. Furthermore, mPFC^SST^-cKO mice compared to WT controls did not exhibit altered exploration or center time in the open field test, open arm time in the elevated plus maze, or time in a lit compartment in the light-dark box (Fig. S1H-L), consistent with findings that enhanced freezing to the CS^−^ is not due to generalized increases in threat reactivity. Further, mPFC^SST^-cKO in the primary motor or auditory cortex did not modify discrimination during threat recall day (Fig. 1F,G), demonstrating that SST neuropeptidergic transmission in all cortical regions does not influence cued-threat discrimination in a similar manner.

**Figure 1:**
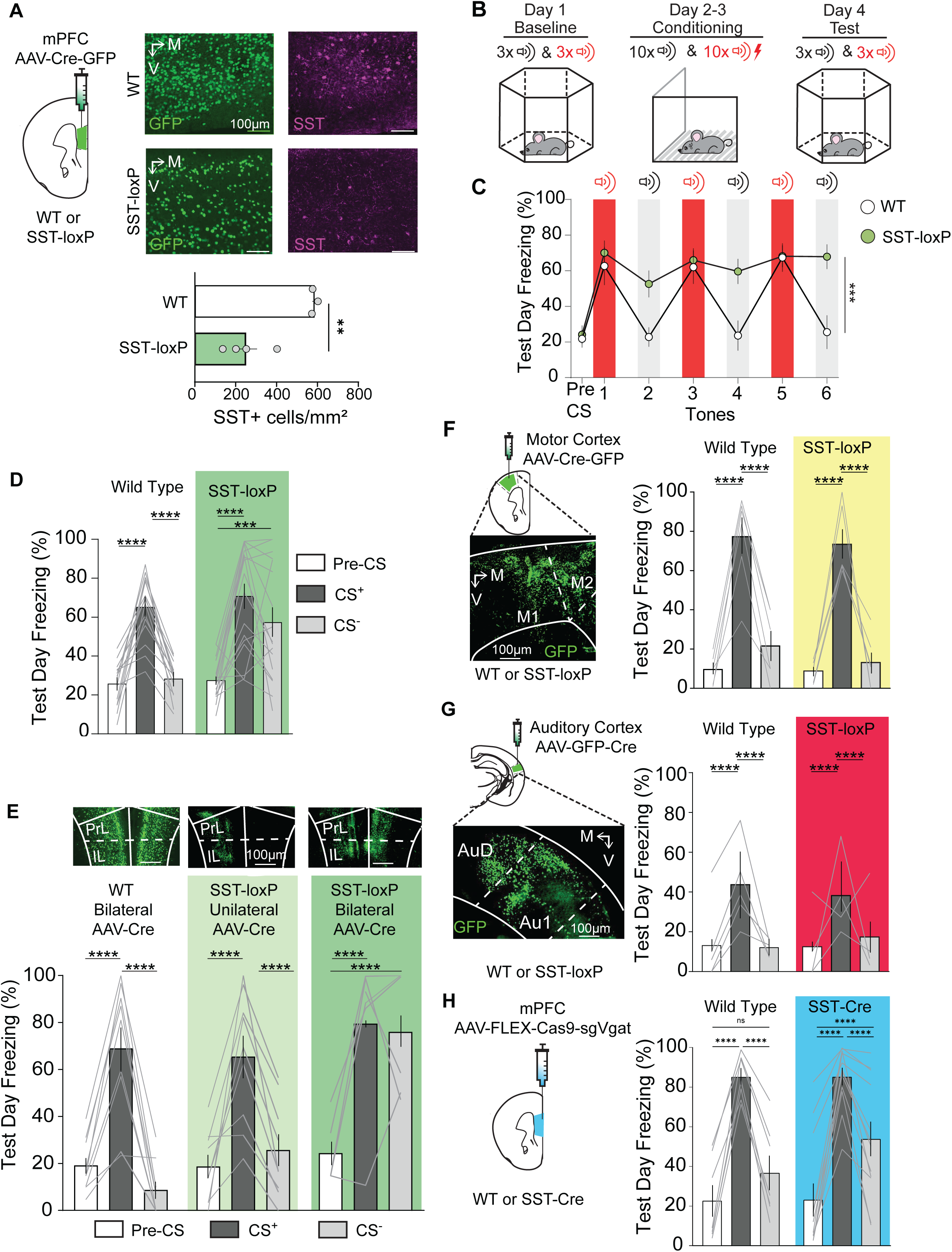
mPFC^SST^ knockdown impairs cued auditory threat discrimination. (A) Bilateral injections of AAV-Syn-Cre-GFP in WT or SST-loxP mice for mPFC^SST^-cKO. Expression of Cre-GFP (green) and SST-immunoreactivity (magenta) in WT (top row) and SST-loxP mice (bottom row). Density of mPFC^SST^-positive cells (Unpaired t-test, \*\**p*=0.0042). (B) Cued threat discrimination timeline. Baseline session where CS^+^ and CS^−^ are presented without consequence (Context A). Conditioning sessions wherein a footshock is paired with the CS^+^, while a CS^−^ signals a neutral outcome (2 days, Context B). Test day where CS^+^ and CS^−^ are presented to elicit cued threat and neutral memories, respectively (Context A). (C) Test day freezing during the pre-CS period, and all subsequent CS^+^ (red shading), and CS^−^ presentations (grey shading; Two-way ANOVA, Genotype x Time Interaction, \*\*\**p*=0.0023). (D) Mean freezing during the pre-CS period, CS^+^, and CS^−^ (Two-way ANOVA, Genotype x Trial Interaction, \*\*\**p*=0.0067; Tukey’s Post-hoc test \*\*\*\**p*<0.0001 WT Pre-CS vs WT CS^+^, \*\*\*\**p*<0.0001 WT CS^+^ vs WT CS^−^, \*\*\*\**p*<0.0001 SST-loxP pre-CS vs SST-loxP CS^+^, \*\*\**p*<0.0001 SST-loxP pre-CS vs SST-loxP CS^−^). (E) Representative images of bilateral injections of AAV-Syn-Cre-GFP in WT or unilateral or bilateral injections in SST-loxP mice. Freezing during threat discrimination recall (Two-way ANOVA, Genotype x Trial Interaction, \*\*\**p*=0.0002; Tukey’s Post Hoc test, \*\*\*\**p*<0.0001 WT Pre-CS vs WT CS^+^, \*\*\*\**p*<0.0001 WT CS^+^ vs WT CS^−^, \*\*\*\**p*<0.0001 unilateral SST-loxP pre-CS vs unilateral SST-loxP CS^+^,\*\*\*\**p*<0.0001 unilateral SST-loxP CS^+^ vs unilateral SST-loxP CS^−^, \*\*\*\**p*<0.0001 bilateral SST-loxP Pre-CS vs bilateral SST-loxP CS^+^, \*\*\*\**p*<0.0001 bilateral SST-loxP Pre-CS vs bilateral SST-loxP CS^−^). (F) Bilateral injections of AAV-Syn-Cre-GFP in WT or SST-loxP mice in the motor cortex and representative image of Cre-GFP expression (Two-way ANOVA, Genotype x Trial Interaction, \*\*\**p*=0.7177; Tukey’s Post Hoc test, \*\*\**p*<0.0001 WT Pre-CS vs WT CS^+^, \*\*\**p*<0.0001 WT CS^+^ vs WT CS^−^, \*\*\**p*<0.0001 SST-loxP Pre-CS vs SST-loxP CS^+^, \*\*\**p*<0.0001 SST-loxP CS^+^ vs SST-loxP CS^−^). (G) Schematic depicting bilateral injections of AAV-Syn-Cre-GFP in WT or SST-loxP mice in the auditory cortex and representative image of Cre-GFP expression. Mean freezing during the test day (right; Two-way ANOVA, Genotype x Trial Interaction, *p*=0.9675; Tukey’s Post Hoc test, \*\*\*\**p*<0.0001 WT Pre-CS vs WT CS^+^, \*\*\*\**p*<0.0001 WT CS^+^ vs WT CS^−^, \*\*\*\**p*<0.0001 SST-loxP Pre-CS vs SST-loxP CS^+^, \*\*\*\**p*<0.0001 SST-loxP CS^+^ vs SST-loxP CS^−^). (H) CRISPR-based strategy to selectively knockout vGAT expression in mPFC^SST^ interneurons. Cued discrimination at cue recall in controls and mice with mPFC^SST^ interneuron vGAT-cKO (Two-way ANOVA, Genotype x Trial Interaction, *p*=0.1666; Tukey’s Post Hoc test, \*\*\**p*<0.0001 WT Pre-CS vs WT CS^+^, \*\*\**p*<0.0001 WT CS^+^ vs WT CS^−^, \*\*\**p*<0.0001 vGAT-cKO Pre-CS vs vGAT-cKO CS^+^,\*\*\**p*<0.0001 vGAT-cKO pre CS vs vGAT-cKO CS^−^, \*\*\**p*<0.0001 vGAT-cKO CS^+^ vs vGAT-cKO CS^−^).

We determined whether mPFC^SST^-cKO mice may alter aversive or nociceptive responses to noxious stimuli that could drive enhanced generalized defensive behavior to neutral cues. However, mPFC^SST^-cKO and control mice displayed similar paw withdrawal latencies in both hot and cold plate assays, and similar thresholds to mechanical stimuli (von Frey assay; Fig. S1M-P), demonstrating that nociceptive defensive behaviors are not modified by mPFC^SST^ ablation.

Because interneuron that release SST also co-release GABA, we compared the impact of genetically suppressing GABAergic transmission from SST interneurons, using Cre-dependent CRISPR knockdown of vesicular GABA transporter (vGAT) (Hunker et al., 2020; Soden et al., 2023) in SST-Cre mice (Fig. 1H). In contrast to mPFC^SST^-cKO, ablation of vGAT from SST-positive neurons resulted in a minor deficit in cued-threat discrimination (Fig. 1H; Fig. S1P). Collectively, these findings indicate that mPFC^SST^ peptidergic transmission regulates cued threat discrimination learning.

### mPFC^SST^ neuropeptidergic transmission is necessary for discrimination between neutral and appetitive outcomes

To identify whether mPFC^SST^ signaling might regulate discrimination between stimuli predictive of salient and neutral outcomes, we compared control WT mice with SST-loxP mice injected with Cre-expressing AAV (Fig. 2A). As expected, in the novel object recognition test, WT-Cre-GFP mice displayed no preference for two identical objects during the pre-test, but increased preference for a novel object that replaced one of the familiar objects during subsequent discrimination testing (Fig. 2B). However, mPFC^SST^-cKO mice spent similar amounts of time interacting with the familiar and novel objects during the discrimination test, suggesting that observed outcomes in the threat discrimination and novel object tasks may stem from discrimination impairments. Viral-mediated knockdown of vGAT expression in SST interneurons produced similar deficits in novel object recognition as mPFC^SST^-cKO (Fig. 2C).

**Figure 2:**
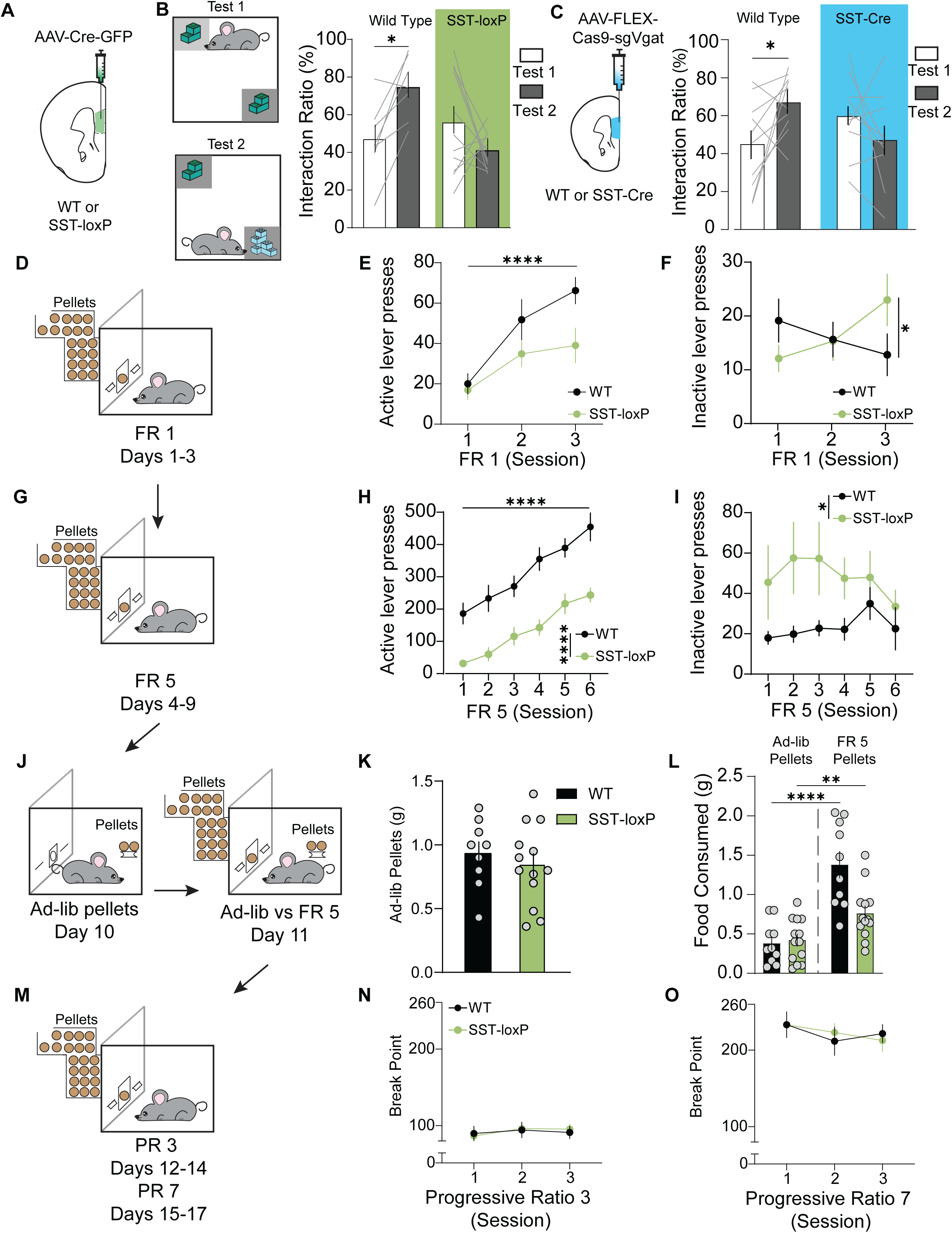
mPFC^SST^ neuropeptidergic regulation of discrimination learning of novel and appetitive outcomes. (A) Schematic of intra-mPFC bilateral injections of AAV-Syn-GFP-Cre onto WT or mPFC^SST^-cKO mice. (B) Experimental timeline for novel object recognition test. Interaction ratio during novel object test 1 and test 2 in WT and SST-loxP mice (Two-way ANOVA, Manipulation x Group Interaction, \*\**p*=0.008; Bonferroni Post-hoc test \**p=*0.0460). (C) Interaction ratio during novel object test 1 and test 2 in WT control and mPFC^SST^ vGAT-cKO mice (Two-way ANOVA, Manipulation x Group Interaction, \**p*=0.0104; Bonferroni Post-hoc test \**p=*0.032). (D,G,J,M) Experimental timeline for operant procedures wherein mice underwent testing under FR1 *(D)* and FR5 *(G)* schedules of reinforcement, forced ad-libitum and ad-libitum vs FR5 choice testing *(J)*, and progressive ratio (PR) testing *(M)*. (E,F) Active *(E;* Two-way ANOVA, Day Main Effect *p*<0.0001) and inactive *(F;* Two-way ANOVA, Day x Genotype Interaction *p*=0.0927) lever presses during FR1 in WT and mPFC^SST^-cKO mice. (H, I) Active (*H*; Two-way ANOVA, Day Main Effect \*\*\*\**p*<0.0001, genotype main effect \*\*\*\**p*<0.0001) and inactive (*I*; Two-way ANOVA, Genotype Main Effect \**p*=0.0425) lever presses during FR5. (K) Ad-libitum chocolate pellet intake in grams (g) in operant chambers in the absence of the active and inactive levers (Unpaired t-test *p*=0.5018). (L) Choice test ad-libitum and FR5 pellet intake in grams (g; One-way ANOVA with Tukey’s Post Hoc test, \*\*\*\**p*<0.0001 WT Ad-libitum vs WT FR5, \*\**p*<0.0052 SST-loxP Ad-libitum vs SST-loxP FR5). (N,O) Breakpoint (total lever presses) during PR3 (*O*; Two Way ANOVA, genotype main effect, *p*=0.9416) and PR7 sessions (*P*; Two-way ANOVA, Genotype Main Effect, *p*=0.8328).

To explore the role of SST signaling in regulating discrimination of appetitive and neutral outcomes, mice were subjected to operant conditioning, in which they had to press an active lever to obtain sucrose pellets, or an inactive lever associated with no outcome (Fig. 2D). WT controls increased active lever presses and decreased inactive lever presses under a fixed ratio 1 (FR1) reinforcement schedule wherein one active lever press resulted in the delivery of one chocolate-sucrose pellet (Fig. 2E,F). In contrast, mPFC^SST^-cKO mice increased both active and inactive lever presses under an FR1 schedule, indicative of deficits in discriminating between the active and inactive lever (Fig. 2E,F; Fig. S2A). We subsequently increased lever press requirements to obtain reward using an FR5 reinforcement schedule (Fig. 2G). Switching to an FR5 results in unrewarded lever presses on the active lever, making it harder to discriminate between outcomes associated with active and inactive lever pressing. Control mice maintained similar levels of discrimination between the active lever and inactive lever on an FR5 schedule relative to FR1 (Fig. 2H, I). mPFC^SST^-cKO mice had increased inactive lever pressing and reduced active lever pressing relative to controls. The deficit in accuracy under the FR5 reinforcement schedule remained for four days of the FR5 schedule before rates of discrimination converged with controls during the last two days of FR5 (Fig. S2B). These results provide further evidence that mPFC^SST^ signaling may facilitate discrimination between actions at active and inactive levers that lead to salient and inconsequential outcomes, respectively.

Next, we examined whether deficits in active lever pressing or enhanced inactive lever pressing found in mPFC^SST^-cKO might be mediated by deficits in motivation. No differences in *ad-libitum* sucrose pellet intake or laboratory chow were observed when mice were given access to the operant box in the absence of operant-derived sucrose (Fig 2K; Fig. S2C). Furthermore, mPFC^SST^-cKO and control mice consumed similar quantities of *ad-libitum* sucrose pellets during simultaneous access to lever press-derived sucrose pellets and freely available sucrose pellets (Fig.2L). WT and mPFC^SST^-cKO mice exhibited similar breakpoints during progressive ratio testing (Fig. 2N,O), consistent with intact motivation with mPFC^SST^-cKO. Moreover, mPFC^SST^-cKO had decreased sucrose preference relative to controls (Fig. S2E). In conjunction with operant studies demonstrating intact reward-seeking behavior when sucrose is freely available, these results provide further evidence that mice fail to appropriately discriminate between the sucrose– and water-containing bottles. Together, these results demonstrate that mPFC^SST^ signaling is critical for discrimination between neutral and salient outcomes, irrespective of valence.

### mPFC^SST^ interneurons are activated during threat discrimination

If mPFC^SST^ transmission is necessary for threat discrimination, then SST interneurons may form representations underlying associative learning. To determine activity of mPFC^SST^ interneurons during associative aversive learning, we performed single-cell Ca^2+^ imaging of mPFC^SST^ interneurons using miniaturized head-mounted microscopes (Fig. 3A). Modulation of single-cell Ca^2+^ activity by speed and distinct task features, including footshock, CS^+^, CS^−^, the intertrial interval following a CS^+^ trial (ITI^+^) or a CS^−^trial (ITI^−^), was assessed using a multiple linear regression encoder model with a temporal kernel (Fig. 3B). During cued threat discrimination conditioning, the majority of mPFC^SST^ neurons responded to footshocks (Fig. 3C,D). Approximately 45% of mPFC^SST^ neurons were modulated by initial CS^+^ and CS^−^ tone presentations during cued threat conditioning acquisition, with the percentage of CS^+^ and CS^−^ modulated neurons decreasing within the session (Fig. 3E). This rapid adaptation in responding prompted us to examine whether cells that retained CS modulation late in threat conditioning sessions were the same neurons modulated early in the session. Cumulative probability of CS^+^ and CS^−^ modulated neurons demonstrated that the majority of neurons were modulated during early trials (Fig. 3F,G). Moreover, CS modulated neurons activated in later trials had a low probability of being modulated *de-novo* and were modulated by earlier tone presentations (Fig. 3H). CS^+^ and CS^−^ triggered Ca^2+^ responses were similar (Fig. 3I) but differed in response at footshock or neutral outcomes, respectively. Further, neurons modulated during later CS^+^ presentations in the session (late CS^+^) displayed CS^+^ evoked activity shifted towards the footshock period and enhanced Ca^2+^ responses at footshock outcome (Fig. 3J), suggesting that mPFC^SST^ interneuron cell assemblies with stable CS encoding associate threat outcomes and threat-predictive cues. The percentage of neurons whose activity was modulated by the intertrial intervals following CS^+^ (ITI^+^) and CS^−^ (ITI^−^) also decreased across the conditioning session (Fig. 3K). Interestingly, ITI^+^ and ITI^−^ modulated neurons displayed sustained Ca^2+^ activity that persisted well beyond footshock (beyond GCaMP kinetics) and at CS^−^ offset, respectively (Fig. 3L), suggesting that mPFC^SST^ interneurons also represent information regarding time periods following threat and neutral outcomes.

**Figure 3:**
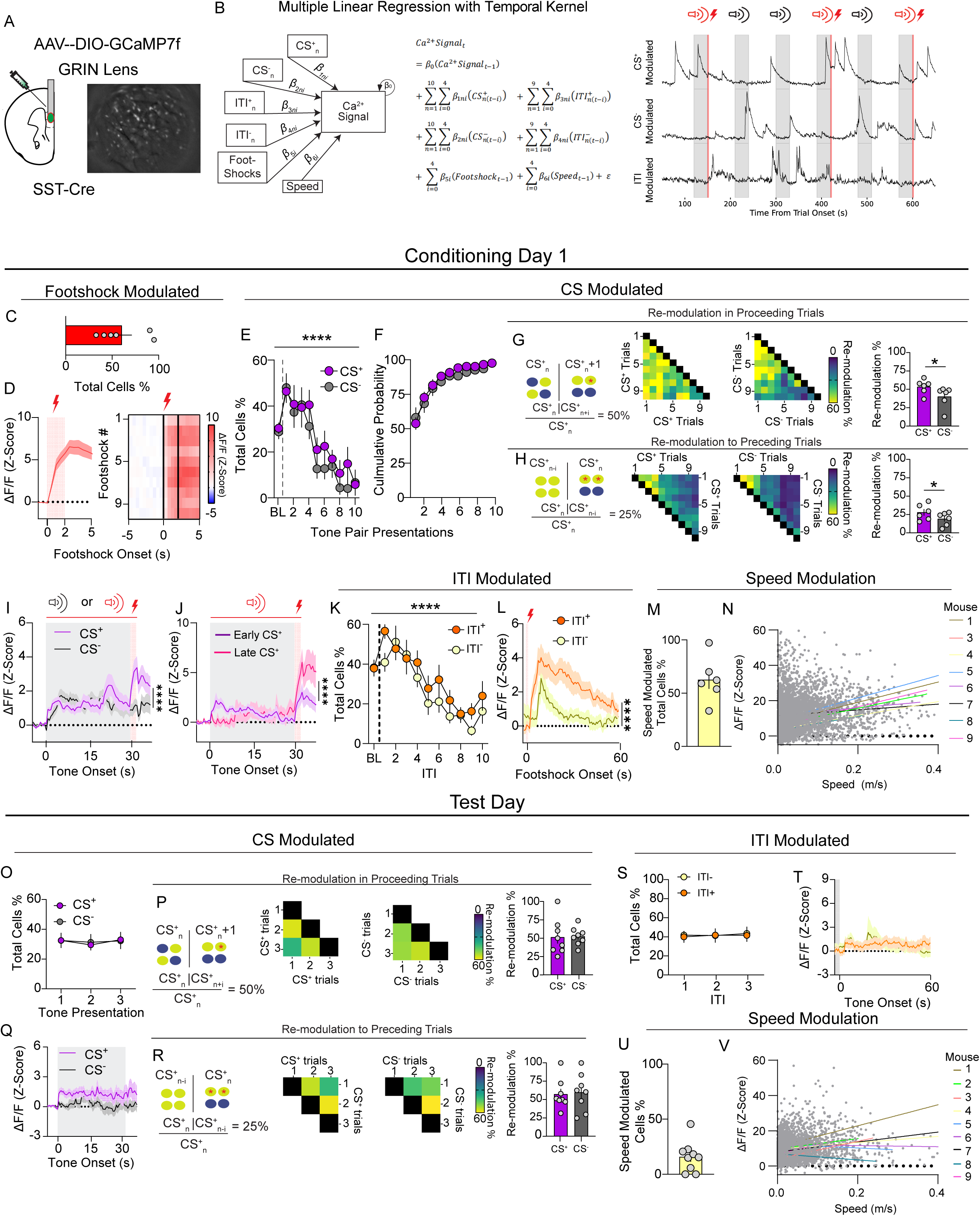
mPFC^SST^ interneurons encode footshock outcomes, neutral and threat-predictive cues, and post-threat periods. (A) GRIN lens implantation and GCaMP recordings from mPFC SST interneurons from SST-Cre mice. (B) Multiple linear regression encoder model with a temporal kernel used to identify neurons whose activity is significantly modulated by task features and speed. Representative SST neurons whose activity is modulated by CS^+^, CS^−^, and ITI. Grey and red highlighted sections of the trace represent CS presentation and footshock delivery, respectively. (C) Percentage of mPFC^SST^ neurons modulated by footshocks during conditioning day 1. (D) Z-Scored GCaMP responses aligned to footshocks collapsed across trials and heatmap of Ca^2+^ responses across trials during conditioning day 1. (E,O) CS modulation of mPFC^SST^ neuron activity across CS^+^ and CS^−^ tone presentations during conditioning day 1 session (E; Two-way ANOVA, Time Main Effect \*\*\*\**p*<0.0001) and test day (O; Two-way ANOVA, Time Main Effect *p*=0.8668). (F) Cumulative probability of CS modulated neurons demonstrating that most CS modulated neurons are recruited in early trials, with no difference between CS^+^ and CS^−^ (Two-way ANOVA, Stimulus Main Effect *p=*0.5110). (G,P) Percentage of neurons whose activity is modulated during CS_n_ and also subsequently modulated in proceeding trials CS_n+I_ during conditioning day 1 (G) and test day (P). Mean percentage of neurons modulated in subsequent trials in conditioning day 1 (G; Paired t-test, \**p*=0.0169) and test day (P; *p*=0.9101). (H,R) Same as G,P but for neurons whose activity was modulated during CS_n_ and during preceding trials CS_n-I_ for conditioning day 1 (H; Paired t-test, \**p*=0.0136) and test day (R; *p*=0.6981). (I,Q) Mean Z-Scored Ca^2+^ activity in CS modulated neurons across all trials during conditioning day 1 (*I;* Two-way ANOVA, Time x Stimulus Interaction \*\*\*\**p*<0.0001) and test day (*Q;* Two-way ANOVA, Time Main Effect *p*=0.2078). (J) Z-Scored Ca^2+^ activity in SST interneurons whose activity was modulated during early CS^+^ (trials 1-3) and late CS^+^ (trials 8-10; Two-way ANOVA, Time x Stage Interaction \*\*\*\**p*<0.0001). (K,S) Percentage of neurons whose activity was modulated during the ITI following a CS^+^ (ITI^+^) and CS^−^ (ITI^−^) (K; Two-way ANOVA, Time Main Effect \*\*\*\**p*<0.0001, ITI Type Main Effect *p*=0.2798) (S; Two-way ANOVA, Time Main Effect *p*=0.9615, ITI Type Main Effect *p*=0.8445). (L,T) Ca^2+^ activity in ITI^+^ and ITI^−^ modulated neurons collapsed across trials during conditioning day 1 (*L*; Two-way ANOVA, Time x ITI Type Interaction \*\*\*\**p*<0.0001) and the test day (*T*; Two-way ANOVA, Time x ITI Type Interaction *p*=0.9216). (M,U) Percentage of neurons modulated by speed. (N,V) Speed modulated neurons did not respond to changes in speed in a linear manner during the conditioning day 1 (N; See statistics table for Pearson Correlation coefficients and *p* values for individual mice) and test day (V).

Similar modulation of mPFC^SST^ interneuron Ca^2+^ activity was present on day 2 of threat conditioning procedures with the exception that ITI^+^ activity was enhanced relative to day 1, with no change in ITI^−^ (Fig. 3L, Fig. S3R). In contrast to the baseline session where mPFC^SST^ interneurons responded to the onset of the CS^+^ and CS^−^ sensory stimuli (Fig. S3A), mPFC^SST^ interneurons displayed sustained activity during the entire CS^+^ period during the test day when mice expressed learned cued threat discrimination (Fig. 3Q), consistent with a shift to threat outcome prediction after associative learning. In contrast, the percentage of mPFC^SST^ interneurons activated by cues, ITI period, or cue modulation in proceeding or preceding trials was similar in baseline and test sessions (Fig. 3O-S; Fig. S3A-E). These results suggest that heightened mPFC^SST^ interneuron activity in periods preceeding and following outcomes linked to specific cues during associative learning may encode information relevant to discrimination learning.

The percentage of mPFC^SST^ interneurons modulated by speed was increased during conditioning days (Fig. 3M, Fig. S3S) and the test day after conditioning (Fig. 3U) relative to the baseline CS exposure session (Fig. S3G). Ca^2+^ activity differed between immobile and mobile states, but was not linearly related to speed (Fig. 3N,V; Fig. S3H,T), despite the percentage of neurons modulated by speed increasing with associative learning and threat discrimination retrieval during the test day. Thus, it is possible that mPFC^SST^ interneurons may respond to defensive state, which under these testing conditions manifest as changes in locomotion (freezing). Together, these results provide a basis wherein mPFC^SST^ interneurons may facilitate associative learning by encoding multiple relevant task variables and footshock outcomes.

### mPFC^SST^ release during cued auditory threat discrimination

SST may be released in the mPFC and potentially activate cognate SST G-protein coupled receptors in response to mPFC^SST^ interneuron activity during aversive associate learning to shape discrimination between aversive and neutral outcomes. However, due to technical constraints, mPFC^SST^ peptide release dynamics have yet to be explored. To address this gap, we utilized fiber photometry recordings using a novel genetically-encoded fluorescent sensor based on an SST receptor (GRAB_SST_) to track SST-peptide dynamics during behavior (Wang et al., 2023 Fig. 4A). We first validated the GRAB_SST_ sensor in WT mice with intra-mPFC injection of GRAB_SST_-expressing virus using *ex-vivo* acute brain slice photometry (Fig. 4A). We found dose-dependent GRAB_SST_ responses evoked by SST bath application, indicating that GRAB_SST_ can reliably detect extracellular SST. We subsequently injected AAV-GRAB_SST_ into the mPFC of WT mice to assess putative SST release during cued threat discrimination acquisition and expression using *in-vivo* fiber photometry (Fig 4B). We observed GRAB_SST_ responses evoked by footshock delivery that persisted into the ITI period during the first day of cued threat conditioning (Fig. 4C), with attenuated footshock responses on conditioning day 2 (Fig. S4B). Since the quantity and intensity of shocks are identical on the first and second day of conditioning, reduced GRAB_SST_ responses suggest SST release is not driven by the footshock outcome per se. GRAB_SST_ responses were not observed in response to the CS^+^ during habituation (before conditioning with the footshock) or the neutral CS^−^ cue during habituation (Fig. S4A) or conditioning (Fig. 4C), indicating that neutral outcomes fail to trigger SST peptide release. Further, mPFC^SST^ interneurons show robust Ca^2+^ responses to footshocks on day 2 of conditioning (Fig. S3I), during the ITI^−^ period following CS^−^ offset (Fig. 3L; Fig. S3R), and in response to the CS^+^ with learning (Fig. 3I,J,Q; Fig. S3O,P), but not reliable GRAB_SST_ responses during these specific time periods. These results suggest there is a dissociation between mPFC^SST^ neuron activation and neuropeptide release. Lastly, we observed that neither the CS^+^ nor CS^−^ evoked GRAB_SST_ responses during the cued threat recall day (Fig. 4D), consistent with time-dependent SST release during associative learning.

**Figure 4:**
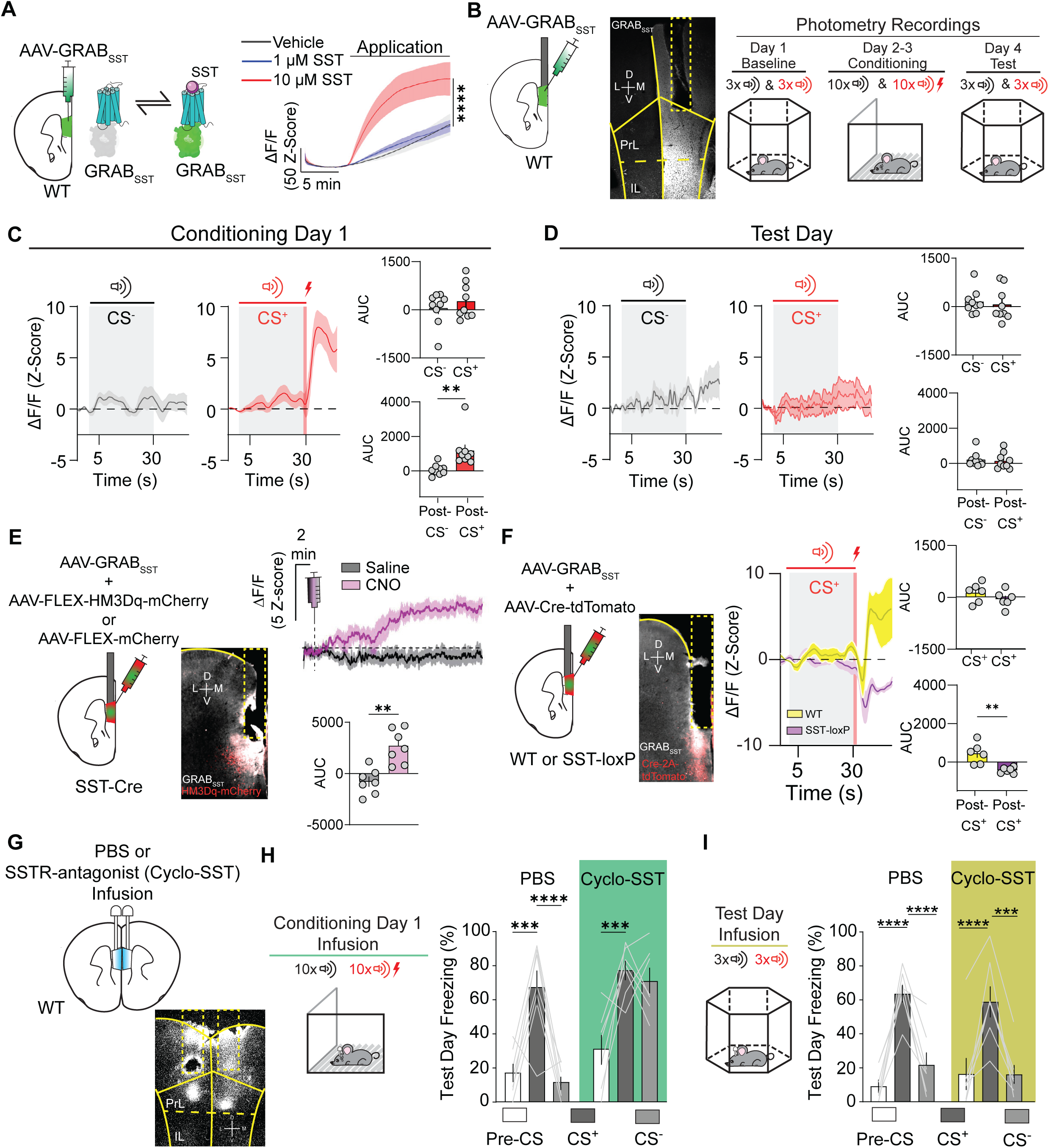
mPFC SST release during cued auditory threat discrimination promotes discriminative learning. (A) Schematic of AAV9-hSyn-GRAB_SST2.0_ (AAV-GRAB_SST_) injection in the mPFC of WT mice. SST application increases GRAB_SST_ fluorescence recorded using photometric recordings in acute *ex-vivo* brain slices in a dose-dependent manner (Two-way ANOVA, Time x Dose Interaction \*\*\*\**p*<0.0001). (B) Schematic depicting *in-vivo* fiber photometry recordings in mPFC AAV-GRAB_SST_-expressing mice during the cued threat discrimination task. (C) GRAB_SST_-mediated fluorescence during presentation of the CS^−^ (left) and CS^+^ (right) during threat discrimination acquisition. AUC of the time windows for the CS^+^ and CS^−^ (top; Paired t-test *p=*0.4052) and the footshock period (bottom; Paired t-test \*\**p*=0.0049). (D) Same as in C but during threat discrimination recall test. AUC of the time windows for the CS^+^ and CS^−^ (top; Paired t-test *p*=0.6804) and post-CS periods (bottom; Paired t-test *p*=0.7328). (E) Schematic demonstrating approach to chemogenetically activate mPFC^SST^ interneurons and monitor GRAB_SST_ fluorescence with *in-vivo* fiber photometry. Systemic administration of CNO (5 mg/kg; i.p.), but not saline, increased GRAB_SST_ fluorescence (top). AUC of GRAB_SST_ responses after saline and CNO (bottom; Paired t-test \*\**p*=0.0011). (F) Schematic depicting strategy to monitor GRAB_SST_ fluorescence with *in-vivo* fiber photometry in WT controls and mPFC^SST^-cKO mice. Time course of cue– and footshock-evoked GRAB_SST_ responses in WT control and mPFC^SST^-cKO mice during threat discrimination. AUC of the time windows for the CS^+^ and CS^−^ (top; Paired t-test *p*=0.2502) and the footshock period (bottom; Paired t-test \*\**p*=0.0033). (G) Schematic depicting implantation of WT mice with bilateral guide cannula for microinjection of the SST receptor antagonist Cyclo-SST. (H) Freezing during recall in mice that were microinjected with vehicle (PBS) or Cyclo-SST during conditioning day 1 (Two-way ANOVA, Treatment x Trial Interaction, \*\**p*=0.0046; Tukey’s Post-hoc test \*\*\**p*=0.0002 PBS Pre-CS vs PBS CS^+^, \*\*\*\**p*<0.0001 PBS CS^+^ vs PBS CS^−^, \*\*\**p*=0.0006 Cyclo-SST pre-CS vs Cyclo-SST CS^+^, *p*=0.8169 Cyclo-SST CS^+^ vs Cyclo-SST CS^−^). (I) Freezing in mice microinjected with vehicle (PBS) or Cyclo-SST prior to the test recall session (Two-way ANOVA, Treatment x Trial Interaction, *p*=0.4054; Tukey’s Post-hoc test \*\*\*\**p*<0.0001 PBS Pre-CS vs PBS CS^+^, \*\*\*\**p*<0.0001 PBS CS^+^ vs PBS CS^−^, \*\*\**p*<0.0001 Cyclo-SST pre-CS vs Cyclo-SST CS^+^, \*\*\**p*<0.0001 Cyclo-SST CS^+^ vs Cyclo-SST CS^−^).

To determine whether GRAB_SST_ tracked endogenous SST release, we injected SST-Cre mice with AAV-expressing the excitatory chemogenetic actuator, HM3Dq-mCherry, or a control vector expressing mCherry in addition to AAV-GRAB_SST_ in the mPFC (Fig. 4E). Chemogenetic activation of mPFC^SST^ interneurons using Clozapine N-oxide (CNO) increased mPFC GRAB_SST_ fluorescence (Fig. 4E), an effect not observed in CNO-treated mCherry controls and both groups when injected with saline (Fig. 4E, Fig. S4C). Importantly, mCherry-expressing control mice displayed footshock-evoked GRAB_SST_ responses similar to HM3Dq-mCherry mice, demonstrating that the lack of CNO response is not due to inefficient GRAB_SST_ expression and/or fiber placement relative to expression (Fig. S4D). To determine whether observed GRAB_SST_ responses during associative aversive learning are due to SST release, we monitored GRAB_SST_ in mPFC^SST^-cKO and control mice. We observed that in contrast to controls, mPFC^SST^-cKO mice expressing GRAB_SST_ did not display footshock-evoked fluorescence (Fig. 4F). These results demonstrate that GRAB_SST_ fluorescence reflects SST neuropeptide release from SST interneurons, and not extrinsic sources. Overall, our findings suggest that SST release in the mPFC does not track the expression of cued threat discrimination, only its acquisition. These results suggest there is a critical time window during which SST release may modulate associations necessary for CS^+^ and CS^−^ discrimination.

### mPFC^SST^ receptor antagonism during acquisition, but not expression, of cued auditory threat discrimination task drives discrimination deficits

To test the hypothesis that SST signaling during the formation of cue-outcome associations is crucial for subsequent discrimination during expression, WT mice were implanted with bilateral guide cannulae aimed at the mPFC to pharmacologically block SST transmission during acquisition (Fig. 4G,H). On the first day of cued threat discrimination conditioning, mice were infused with vehicle (control) or the SST receptor antagonist, cyclo-SST (Fig. 4H). Interestingly, intra-mPFC SSTR antagonism impaired CS^+^/CS^−^ discrimination characterized by increased freezing behavior on test day in response to the CS^−^, in contrast to vehicle-treated mice (Fig. 4H). This indicates that SST-mediated activation of SSTRs during the acquisition of cue outcome associations is necessary for discrimination during expression. Cyclo-SST-treated mice did not display enhanced freezing during the pre-CS period of the threat expression test (Fig. 4H,I). Importantly, mPFC cyclo-SST infusion during acquisition did not modify freezing elicited by footshocks, in response to the CS^+^ and CS^−^ (Fig. S4E,F), demonstrating that enhanced freezing to the CS^−^ is not a consequence of enhanced reactivity during acquisition. We subsequently determined whether mPFC SSTR antagonism during recall of cued threat discrimination, when no SST release is observed (Fig. 4D), would enhance freezing to the CS^−^. Vehicle controls and intra-mPFC cyclo-SST-treated mice displayed similar discrimination between the CS^+^ and CS^−^ (Fig. 4I). Together with our *in-vivo* GRAB_SST_ recordings, these results suggest that SST release and subsequent SSTR activation during learning is important for establishing that footshock-predictive and neutral cues are salient and inconsequential, respectively, a process essential for appropriate behavioral discrimination in other settings.

### SST knockdown impacts mPFC dynamics during cued auditory threat discrimination learning

It is possible that SST signaling may shape mPFC dynamics during the formation of cue-outcome associations essential for discriminating between neutral and salient outcomes. To determine how selective SST ablation in the mPFC impacts circuit dynamics during cued threat discrimination we used microendoscopic single-cell Ca^2+^ imaging of pan-neuronal mPFC cells in freely-moving WT and SST-loxP mice injected with a cocktail of AAV-Cre-tdTomato and AAV-GCaMP7f (Fig. 5A). A multiple linear regression with a temporal kernel was used to identify task and behaviorally-relevant features that predicted Ca^2+^ activity in each neuron (Fig. 3B; Fig. 5B; Wang et al., 2024). This model uncovered no differences in the percentage of neurons whose activity was modulated by either the CS^+^ or CS^−^ (Fig. S5A,C), ITI (Fig. S5M), or speed (Fig. S5N) during baseline habituation. Further, Ca^2+^ responses aligned to these events during the baseline day of cued auditory threat discrimination is comparable between WT and mPFC^SST^-cKO mice (Fig. S5B, D). Interestingly, during the first and second day of cued threat discrimination conditioning, SST-ablation decreased the percentage of neurons encoding footshock and their time locked responses (Fig. 5C,D; Fig.S6O,P). The percentage of cells encoding the CS^+^ and CS^−^ increased during conditioning relative to the encoding of cues during the baseline day in control mice, an effect that was absent in mPFC^SST^-cKO mice (Fig. 5E,G; Fig. S5A,C; Fig. S6A,C). During conditioning day 1, Ca^2+^ responses time-locked to CS^+^ onset in CS^+^ modulated neurons were similar in control and mPFC^SST^-cKO mice (Fig. 5F), in contrast to CS^−^ modulated cells, which had significantly smaller Ca^2+^ responses in controls relative to mPFC^SST^-cKO mice (Fig. 5H). Together, these results suggest that mPFC^SST^-cKO mice fail to display heightened Ca^2+^ activity to the CS^+^ relative to the CS^−^ observed in controls (Fig. 5F, H). Further, increased CS activity highlights that decreased Ca^2+^ activity to the footshock in mPFC^SST^-cKO mice is not due to non-specific decreases in Ca^2+^ responses to external stimuli. These results suggest that SST neuropeptide transmission shapes encoding of distinct task variables, including threat-predictive and neutral cues, associated with aversive learning.

**Figure 5:**
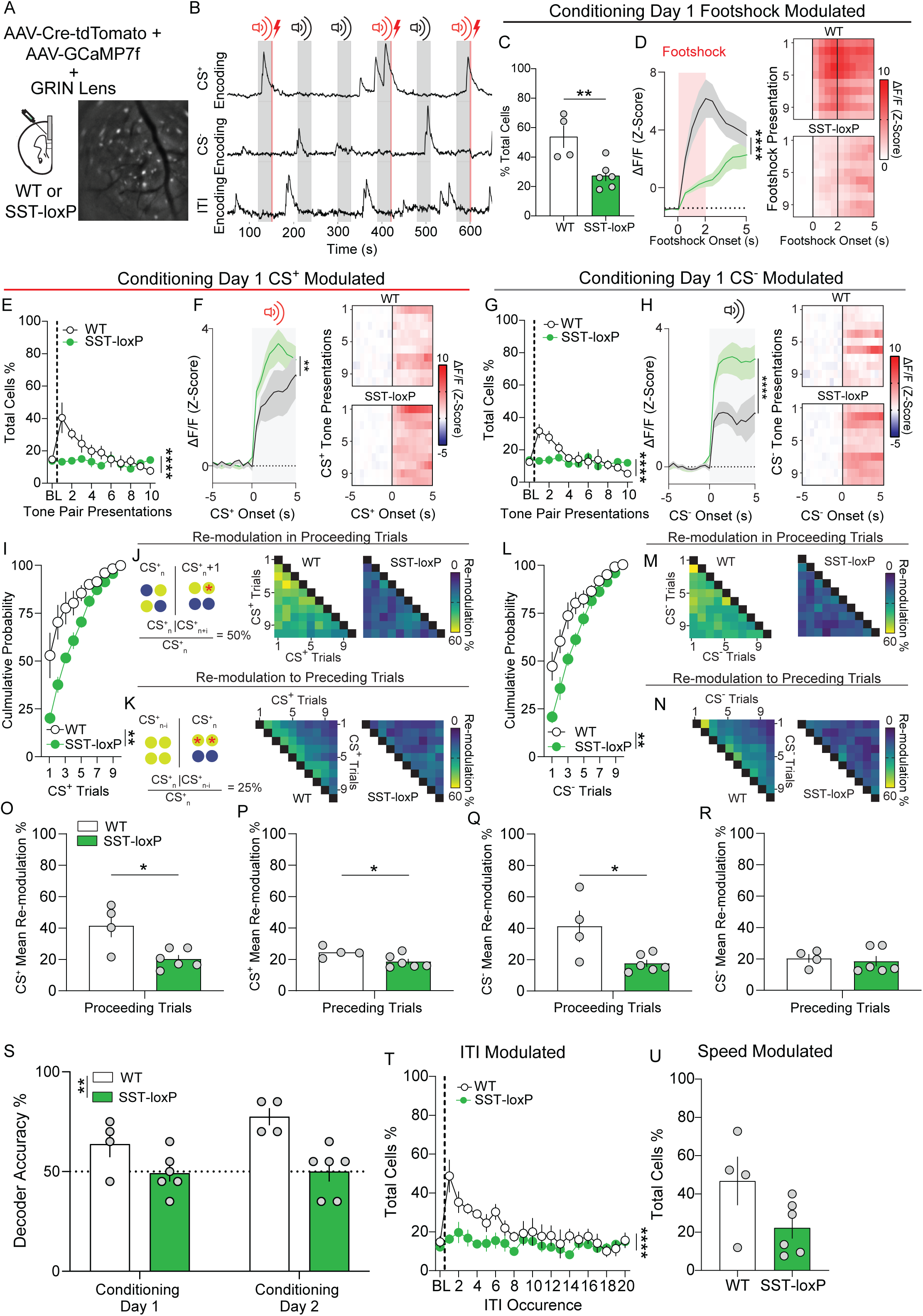
mPFC SST knockdown alters mPFC neuronal encoding in a cued-threat discrimination task. (A) Schematic depicting bilateral injections of AAV9-Syn-jGCaMP7f and AAV5-hSyn-Cre-P2A-tdTomato and GRIN lens implant into the mPFC of WT or SST-loxP mice to record single-cell GCaMP7f activity with or without SST expression, respectively. (B) Representative traces of single neurons encoding CS^+^, CS^−^, or ITI with tones (gray highlighted sections) and footshocks (red highlighted sections) represented. (C,E,G,T,U) Percentage of total neurons encoding footshock (C), CS^+^ (E), CS^−^ (G), ITI (T), and speed (U) during conditioning day 1, WT vs SST-loxP (C, Unpaired t-test, \*\**p*=0.0079; *E*, Two-way ANOVA, Time x Genotype Interaction \*\*\*\**p*<0.0001; G, Two-way ANOVA Time x Genotype Interaction \*\*\*\**p*<0.0001; T, Two-way ANOVA Time x Genotype Interaction \*\*\*\**p*<0.0001; U, Unpaired t-test, *p*=0.0807). (D,F,H) Timecourse of Z-scored GCaMP7f activity of neurons modulated by the footshock (D), CS^+^ (F), and CS^−^ (H), during conditioning day 1. Heatmaps representing Z-scored activity in response to footshocks, CS^+^, and CS^−^ across trials in WT (top) and SST-loxP (bottom) mice. (D; Two-way ANOVA, Time x Genotype Interaction \*\*\*\**p*<0.0001, F; Two-way ANOVA, Time x Genotype Interaction \*\**p*=0.0031, H; Two-way ANOVA Time x Genotype Interaction \*\*\*\**p*<0.0001). (I, L) Cumulative probability of cells that become significantly modulated by the CS^+^ (I) and CS^−^ (L) across conditioning day 1 (I; Two-way ANOVA, Genotype Main Effect \*\**p=*0.0090; L, Two-way ANOVA, Genotype Main Effect \**p=*0.0226). (J,K,M,N) Heatmap representing the percent of neurons modulated to CS^+^ (J,K) or CS^−^ (M,N) tones across conditioning day 1 in WT (left) and SST-loxP (right) neurons. Re-modulation percentage was calculated by determining the percentage of neurons with significant modulation in a specific trial (CS^+^_n_ or CS^−^_n_) that were also significantly modulated in n-proceeding (CS_n+i_) or n-preceding (CS_n-i_) trial (CS_n_ | CS_n±i_). % re-modulation = (CS_n_ | CS_n±i_ / CS_n_). (O, Q) Percentage of neurons with significant modulation in *n*-proceeding trials in WT vs SST-loxP (O, Unpaired t-test \**p*=0.0118; Q, Unpaired t-test \**p*=0.0224). (P, R) Percentage of neurons with significant modulation in *n*-preceding trials in WT vs SST-loxP (P, Unpaired t-test \**p*=0.0455; R, Unpaired t-test \**p*=0.7003). (S) Decoder accuracy is increased in WT relative to SST-loxP mice across two days of conditioning. (Two-way ANOVA, Genotype Main Effect \*\**p=*0.004).

We observed that SST interneurons adapt within conditioning sessions to encode task variables in a sparse subset of interneurons (Fig. 3). SST release during conditioning from mPFC^SST^ interneurons may a play a role in regulating representational adaptation or stability in mPFC neurons. We subsequently investigated whether CS modulated neurons in control and mPFC^SST^-cKO mice displayed differential within-session adaptation or stability in subsets of CS^+^ and CS^−^ encoding cells. CS^+^ and CS^−^ encoding cells were modulated more rapidly during early trials of the conditioning sessions in controls relative to mPFC^SST^-cKO mice (Fig. 5I,L; Fig. S6E,H). To further explore whether CS modulated neurons in controls and mPFC^SST^-cKO mice displayed differential stability or *de-novo* representations across trials, we examined the percentage of neurons whose activity was modulated in past (preceding) or future trials (proceeding). The percentage of neurons modulated in a present trial (CS^+^_n_ or CS^−^_n_) and also modulated in proceeding trials (CS^+^_n+i_ or CS^−^_n+i_) rapidly dropped off across trials in control mice (Fig. 5J,M,O,Q). Conversely, CS^+^ and CS^−^ modulated neurons in the present trial (CS^+^_n_ or CS^−^_n_) had low probability of modulation unless they were modulated in trials immediately preceding the present trial (CS^+^_n-i_ or CS^−^_n-i_) in control mice (Fig. 5K,N,P,R). These results are consistent with the notion a subset of CS modulated neurons form stable, and not *de-novo*, representations. Modulation by CS^+^ and CS^−^ in proceeding, but not preceding, trials was decreased in mPFC^SST^-cKO mice relative to control mice, suggesting that representational adaptation in CS^+^ and CS^−^ encoding is enhanced in mPFC^SST^-cKO mice (Fig. 5J,K,M-R). The percentage of cells with re-modulation to the CS^+^ to proceeding, but not preceding, trials was enhanced in WT mice across conditioning (Fig. S6E-N), but not test day (Fig. S7E-L), relative to baseline (Fig. S5E-L). Together, these results suggest that SST neuropeptidergic transmission facilitates recruitment of stable CS^+^ and CS^−^ representations in subsets of mPFC neurons during associative learning. In agreement with CS modulation of neuronal activity containing SST-dependent threat– and neutral-associated information, decoding of CS^+^ and CS^−^ trials was decreased in mPFC^SST^-cKO mice relative to controls (Fig. 5S). These results suggest that SST neuropeptidergic transmission is necessary for mPFC neuronal discrimination of CS^+^ and CS^−^ trials during acquisition.

The percentage of ITI-modulated neurons was increased during conditioning relative to the baseline and test days in control mice (Fig. 5T, Fig. S5M, Fig. S6Q, Fig. S7M). Increased modulation of mPFC neurons during the ITI period may reflect ongoing evaluation during post-threat and neutral outcome epochs. The increase in the percentage of ITI modulated neurons across learning was not present in mPFC^SST^-cKO mice, suggesting that post-outcome, in addition to cue and footshock, processing is impaired. Speed-modulated cells were not different between the two groups during baseline, conditioning, or test days (Fig 5U; Fig. S5N, Fig. S6R, Fig. S7N). These results suggest that despite regulating freezing to neutral cues, mPFC^SST^-cKO does not impact mPFC neuronal activity associated with changes in movement per se. Together, these results demonstrate that SST neuropeptidergic transmission shapes the encoding of cues predictive of aversive and neutral outcomes and post-outcome periods that may interact during learning.

### mPFC^SST^ signaling modulates configural representations and population level encoding in mPFC associated with threat discrimination

PFC neurons display linear and non-linear mixed selectivity wherein neural encoding of variables can be multiplexed or dependent on task rules, respectively, and coordinated activity at the population-level (Averbeck et al., 2006; Fusi et al., 2016; Panzeri et al., 2022; Tye et al., 2024). SST signaling may not only be influencing how neurons respond to individual stimuli or epochs, but also multiplexing variables, more specifically forming configural or conjunctive representations. In controls, the percentage of CS modulated neurons encoding CS^+^, CS^−^, or both tones within a trial pair is comparable (Fig. 6A,C,D). In contrast, mPFC^SST^-cKO mice show a decrease in the percentage of neurons encoding both the CS^+^ and CS^−^ relative to controls (Fig. 6B,C,D). Interestingly, the percentage of CS modulated neurons encoding the CS^−^ in isolation was enhanced in SST-loxP mice relative to controls (Fig. 6C). This suggests that CS^+^ and CS^−^ multiplexing, but not isolated CS processing, in individual neurons is regulated by SST neuropeptidergic transmission.

**Figure 6:**
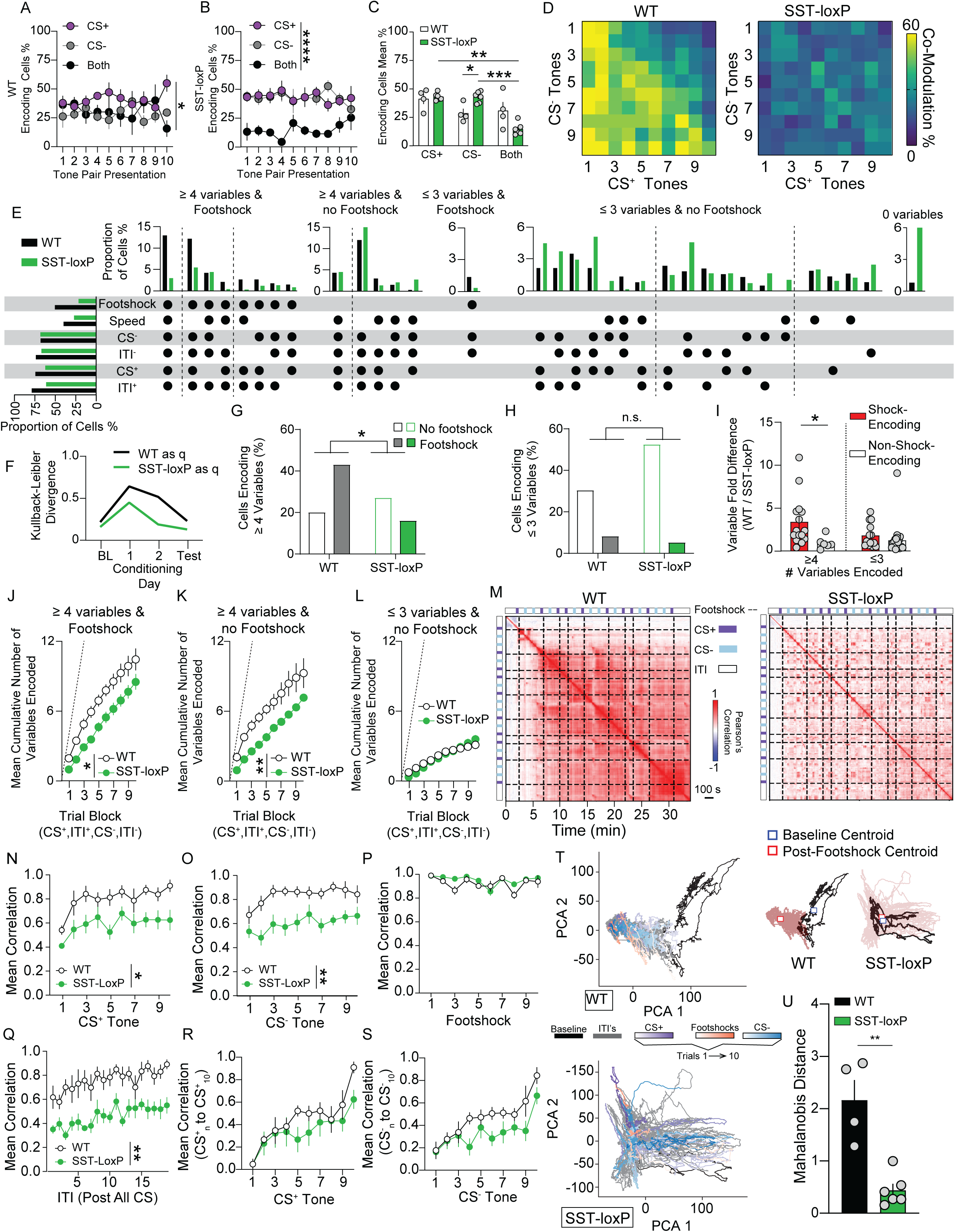
mPFC SST knockdown alters encoding of task variables in a threat discrimination task. (A,B) Percentage of neurons encoding the CS^+^ or CS^−^ individually, or both across CS^+^/CS^−^ trial pairs during acquisition day 1 in WT mice (A; Two-way ANOVA, Time x Encoding Type Interaction \**p*=0.0270) and SST-loxP mice (B, Two-way ANOVA, Encoding Type Main Effect *p*<0.0001). (C) Average individual and joint CS encoding in WT and SST-loxP mice during conditioning day 1 (Two-way ANOVA, Encoding Type x Genotype Interaction, \**p*=0.0103; Tukey’s Post-hoc test, WT vs SST-loxP CS^−^ \**p*=0.0276, SST-loxP CS^+^ vs both \*\*\**p=*0.0006, CS^−^ vs both, \*\**p=*0.0021). (D) Percentage of neurons modulated by the CS^+^ and CS^−^ tone across the discrimination day 1 session in WT (left) and SST-loxP (right) mice. Co-modulation percentage is calculated as the number of neurons with significant modulation to CS^+^ in a given trial and all CS^−^ trials. (E) UpSet visualization of neurons modulated by different combinations of footshock, CS^−^, CS^+^ or ITI following a CS^+^ (ITI^+^) and CS^−^ (ITI^−^). Categories with <1% of neurons modulated are not shown. Left horizontal bars represent percentages of cardinal categories, including those not shown. Dashed lines indicate separation of categories encoding different numbers of variables. (F) Kullback-Leibler divergence between WT and SST-loxP mice as Q (the reference group) across days demonstrating that differences in the probability of combinatorial encoding distributions is smaller during baseline and increases with threat conditioning. (G, H) Percentage of neurons with configural encoding (≥4 variables; G, left; Two-sided Fishers Exact Test *p*=0.0026) or with limited encoding (≤3 variables; H, right; Two-sided Fishers Exact Test *p*=0.1273) including and excluding footshock in WT and SST-loxP mice. (I) The variable fold difference between WT and SST-loxP mice in distinct categories (each represented by a dot) containing (red) or lacking a footshock (gray) with configural encoding (≥4 variables; left) or limited encoding (≤3 variables; right) of variables. Fold difference is the percentage of neurons encoding a distinct category in WT mice divided by the percentage of neurons in SST-loxP mice. Fold differences greater and less than 1 indicate more encoding in WT and SST-loxP mice for a category, respectively (Two-way ANOVA, Variables x Footshock Interaction, *p=*0.1317, Footshock main effect, \**p=*0.0162; Bonferroni multiple comparison \**p=*0.0323). (J-L) Mean cumulative sum during conditioning day 1 of variables encoded across trial blocks consisting of a CS^+^, ITI^+^, CS^−^, and ITI^−^, in cells with configural encoding (≥4 variables) with (J) or without footshock (K), or in cells with limited encoding (≤3 variables) without footshock (L). Dashed line represents theoretical maximum number of variables that can be encoded (J, Two-way ANOVA, Genotype Main Effect \**p*=0.0101; K, Two-way ANOVA, Genotype Main Effect \*\**p*=0.0056; L, Two-way ANOVA, Genotype Main Effect *p*=0.7719). (M) Heatmaps representing the mean Pearson’s correlation coefficients of vectorized population Ca^2+^ activity across all time points during conditioning day 1 for WT (left) and SST-loxP mice (right). Black dashed lines indicate shock onset, purple and grey squares on borders indicate CS^+^ and CS^−^ presentations, respectively. (N-S) Mean Pearson’s correlation of activity around CS^+^ (N), CS-(O), footshock (P), ITI (Q), CS^+^ to final tone (R), and CS^−^ to final tone (S) (N, Two-way ANOVA, Genotype Main Effect \**p=*0.0307; O, Two-way ANOVA, Genotype Main Effect \*\**p=*0.0061; P, Two-way ANOVA, Genotype Main Effect *p*=0.1539; Q, Two-way ANOVA, Genotype Main Effect \*\**p=*0.0034; R, Two-way ANOVA, Genotype Main Effect *p*=0.1539; S, Two-way ANOVA, Genotype Main Effect *p*=0.1558). (T) Representative principal component analysis (PCA) trajectories of WT (top) and SST-loxP (bottom) throughout the entirety of conditioning day 1. Distinct task epochs are indicated by color. Early to late trials are indicated by light to dark color gradients, respectively. CS^+^ presentations in purple, CS^−^ in blue, and footshocks in red, with baseline period in solid black and ITI in solid grey. Centroid distances (top right) of WT (left) and SST-loxP (right), blue square indicates baseline centroid, red square indicates post-footshock centroid, respectively). (U) Mahalanobis Distance of WT and SST-loxP mice (Unpaired t-test, \*\**p=*0.0010).

Most mPFC neurons adapt, with only a sparse population maintaining stability in CS modulated activity across the first conditioning session (Fig. 5). Likewise, it is possible that co-modulation by both CS similarly adapts or retains stability across conditioning. The percentage of neurons activated during either CS^+^ or CS^−^ by a proceeding CS^−^ or CS^+^ trials, respectively, was enhanced relative to neurons modulated by a preceding CS^+^ or CS^−^ in WT mice, but not in SST-loxP mice (Fig. S10B). This suggests that co-modulated neurons maintain a stable representation based on early, but not late, tone presentation encoding in an SST-dependent manner. Conversely, there is a very low chance that a cell will *de-novo* encode the CS^+^ and CS^−^ in a subsequent trial, an SST independent process. Differences in conjunctive CS^+^ and CS^−^ encoding between WT and SST-loxP mice were not present on baseline (Fig. S8A-D, S10A) and conditioning day 2 (Fig. S9A-D, S10C). Together, these results suggest that SST signaling during cue-outcome association formation may facilitate configural representations in mPFC neurons pertaining to both neutral and threat-predictive cues, without modulating encoding of CS^+^ and CS^−^ in isolation.

Selective influence of mPFC^SST^-cKO on modulation of mPFC activity by both the CS^+^ and CS^−^, but not by these cues in isolation, provides a basis for SST neuropeptidergic transmission to differentially regulate mPFC activity depending on the combinatorial composition of variables encoded. To this end, we categorized neurons based on the combination of variables encoded across baseline, conditioning, and test sessions (Fig. 6E; Fig. S8E,M; S9E). The Kullback-Leibler divergence between the distribution of percentages of neurons in distinct encoding categories increased during threat conditioning day 1 relative to the baseline, conditioning day 2, and test sessions (Fig. 6F), suggesting that mPFC neurons differentially encode distinct combinations of variables during threat discrimination acquisition and expression. In WT controls the majority of mPFC neurons were modulated by multiple variables, with 42.58% of neurons forming configural representations (≥4 variables) incorporating footshock and 19.9% of configural representations not modulated by footshock (Fig. 6E,G). These results suggest that mPFC configural neuronal representations incorporating salient threat outcomes are over-represented and may facilitate discrimination between distinct cue-outcome associations. The percentage of neurons encoding multiple variables including footshock (≥4 variables) was significantly decreased in mPFC^SST^-cKO mice (16.4%), while the percentage of neurons encoding multiple variables not including shock was enhanced (26.93%; Fig. 6E,G,I). Further, the percentage of neurons with limited encoding (≤3 variables) modulated by footshock or not did not differ between controls and mPFC^SST^-cKO mice (Fig 6E,H,I). The majority of footshock modulated neurons also encoded ≥4 variables, with sparse neurons encoding ≤3 variables and footshock in both WT and SST-loxP mice (Fig. 6G-I). Configural, but not limited (≤3 variables), encoding increased in WT mice throughout the session as variables were introduced, an effect that was attenuated in SST-loxP mice (Fig. 6, J-L), an effect that was also observed in conditioning day 2 (Fig. S9I-K). These results are consistent with the hypothesis that SST neuropeptidergic transmission regulates higher-order representations in mPFC circuits, potentially helping associate threat outcome with other environmental variables and behavioral state within individual neurons.

Given that configural neural representations incorporating threat outcome and all other epochs of the threat conditioning session (e.g. ITIs and CS) are compromised in mPFC^SST^-cKO mice, the possibility exists that mPFC SST signaling is necessary for correlated population-wide activity throughout the conditioning session (Panzeri et al., 2022). Population activity vectors were correlated at any given time point with all other time points throughout the session (Fig. 6M; Fig. S8F,N; S9L). This permits evaluation of population-wide correlated activity on a moment-by-moment basis (e.g. centered around an individual CS_n_) and between specific variable events across a session (e.g. correlations between a CS_n_ and CS_n±i_; Wang et al., 2024). In control mice, population-wide correlated activity was low during the pre-CS period and increased immediately with delivery of the first footshock on a moment-by-moment basis (along the diagonal of the correlation matrix excluding the main diagonal; Fig. 6M-Q; Fig. S9L-P). In contrast, correlated activity at the population level was decreased in mPFC^SST^-cKO mice both on a moment-by-moment basis, except around footshock (Fig. 6M-Q; Fig. S9L-P). We determined how population level activity on individual cue trials correlates with the first or final CS^+^ or CS^−^ as mice associated cues with outcomes (Fig. 6R,S; Fig. S8G,H,O,P; Fig. S9Q,R; Fig. S10E-L). During conditioning day 1, population activity correlations during the first CS^+^ or CS^−^ and subsequent cues was low in both control and SST-loxP mice and increased in subsequent trials, suggesting that cues later in the session are differentially processed relative to those early in the session (Fig. 6R,S; Fig S10F,J). This is consistent with a general switch in population level encoding following the first CS^+^ or CS^−^ as cues become associated with a novel threat or neutral outcome in both WT and SST-loxP mice. Intact differentiation between initial and following cues may provide a basis for SST-loxP mice to develop appropriate conditioned freezing to the CS^+^, whereas defective configural representations and persistent correlated activity in response to threat outcome may provide a basis for inappropriate discrimination. Lastly, analysis of principal component trajectories of mPFC circuit activity revealed that the initial footshock elicited a shift in the state space of WT mPFC dynamics, an effect that was decreased in SST-loxP mice (Fig. 6T,U; Fig. S9S,T). This indicates that SST transmission is necessary for a state-dependent shift in population-wide mPFC dynamics in response to threat. Collectively, these findings demonstrate that mPFC^SST^ neuropeptidergic transmission is necessary for neurons to multiplex salient footshock outcomes with other variables to form configural representations and facilitates population-level encoding associated with discriminative learning.

## Discussion

Here we describe a role for mPFC^SST^ neuropeptidergic transmission in regulating discrimination between salient and neutral outcomes. We demonstrate that mPFC^SST^ release and SSTR signaling during associative threat learning is necessary for subsequent discrimination. mPFC^SST^ neuropeptidergic transmission modulates how mPFC neurons associate salient outcomes to on-going behavioral and environmental state. Collectively, this study provides evidence of endogenous mPFC^SST^ signaling in shaping learning and emergent properties of mPFC circuits, highlighting the importance of a neuropeptidergic signal classically used as a marker of inhibitory interneurons.

mPFC^SST^ interneurons are implicated in threat regulation, reward, and cognition (Abbas et al., 2018; Cummings et al., 2022; Cummings & Clem, 2020; Dao et al., 2021; Joffe et al., 2022; Kim et al., 2016). Technical limitations prevented insights to endogenous SST dynamics and the impact on cortical circuit function. As such, existing models of SST interneuron function are based on their ability to release GABA on postsynaptic pyramidal and inhibitory interneurons. Using a combination of novel interdisciplinary approaches, the present work reveals that mPFC^SST^ release and SSTR signaling in response to threats during conditioning, but not during expression, is critical for discrimination of learned outcomes. SST receptors are Gi-coupled GPCRs that inhibit neurotransmitter release and/or hyperpolarize neurons (Casello et al., 2022; Song et al., 2021). These results suggest that SST receptor-mediated long-term synaptic or intrinsic excitability plasticity may underly changes in mPFC activity that is necessary to subsequently use associative learning rules established during learning later in expression. In contrast, mPFC^SST^ interneuron-mediated GABAergic transmission complements the role of SST transmission in regulating threat discrimination since vGAT knockout marginally produced threat discrimination deficits, yet still produced novel object recognition deficits comparable to mPFC^SST^-cKO. Since GABA_A_ receptors are ligand-gated ion channels with distinct spatial cellular distributions and temporal properties relative to SST receptors, uncovering circuit-level influences of SST neuropeptide / SST receptor and GABA / GABA_A_ receptor signaling may elucidate how cortical interneurons use diverse signaling modalities to orchestrate circuit function. Delineating time-dependent recruitment of SST neuropeptidergic signaling adds a new underappreciated dimension for understanding SST interneurons’ impact on information processing.

This study provides evidence that mPFC^SST^ signaling shapes encoding in mPFC neurons, including configural representations binding salient footshocks to other variables and regulating population-level correlated activity and state space. This highlights a novel role for a “cellular marker” in regulating emergent properties of cortical circuits. Mapping causal inference between neural codes or representations to behavior has been elusive in neuroscience, due to complexity of neural dynamics and use of imprecise experimental perturbations (Jazayeri & Afraz, 2017; Langdon et al., 2023; Panzeri et al., 2022). Uncovering time-dependent SST release and receptor signaling during associative learning and interrogating this change with mPFC^SST^-cKO during single-cell recordings, we provide evidence of a physiologically relevant change in the neural manifold that provides causal inference into the relationship between complex neural codes and discriminative learning. Since mPFC neurons in mice with SST cKO retain representations of the CS^+^ and CS^−^ independently, these results suggests that joint CS^+^ and CS^−^ encoding, and not isolated cue processing, is critical for associative discriminative learning. Further, our work suggest that discrimination of discrete cues also relies on representations combining other variables beyond cues, including outcomes and post-outcome epochs conducive for retrospective evaluation. Local infusion of exogenous SST peptide or the SSTR receptor antagonist, Cyclo-SST, in V1 visual cortex improved and impaired visual perception, respectively (Song et al., 2020). Exogenously applied SST increased preferential visual orientation tuning in layer 4/5 V1 neurons in anaesthetized mice (Song et al., 2020). Our results extend this work demonstrating a role for endogenous SST transmission, which in conjunction with previous work suggest that SST neuropeptide transmission in various cortical circuits regulates information processing essential for behavior. However, our control experiments eliminating SST in motor and auditory cortex suggest that SST transmission in distinct cortical and sub-cortical areas may play a specialized role. GRAB_SST_ recordings in the basolateral amygdala revealed increased fluorescence to reward and reward-predictive cues, but no responses to aversive air puff (Wang et al., 2023). Recently, we demonstrated that dynorphin signaling in the ventral mPFC is essential for population level correlations and state transitions upon footshock without modifying encoding of speed or task features during threat conditioning in individual neurons (Wang et al., 2024). Moreover, unlike SST neuropeptide transmission, dynorphin signaling modified freezing during threat acquisition but not during expression. This contrasts with modification of variable encoding and population-level correlated activity and state transitions in mPFC^SST^-cKO mice and impairments in discrimination learning needed for subsequent expression. Divergent influence of SST and dynorphin signaling on threat-associated behavior and neural coding, suggest that distinct neuropeptides may differentially orchestrate emergent properties of cortical circuits and top-down control of threat outcomes. Together, this provides causal inference into complex mPFC neural coding of salient and neutral outcomes essential for top-down control of affective responses.

Impairments in discrimination observed with mPFC^SST^-cKO parallel symptom clusters observed in various mental health disorders. Decreased PFC SST expression and immunoreactivity is documented in depression, bipolar disorder, and schizophrenia, as well as in amygdala and hippocampus (Beneyto et al., 2012; Fung et al., 2014; Hashimoto et al., 2008; Lin & Sibille, 2015; Morris et al., 2008; Pantazopoulos et al., 2017; Seney et al., 2015; Tripp et al., 2011). These disorders are marked by deficits in top-down control required for adaptive behaviors in response to neutral and salient threatening and reward outcomes. This includes negative bias in mood and anxiety disorders consisting of enhancements in aversive processing and mal-adaptive response to innocuous events (Kenwood et al., 2022; Pizzagalli & Roberts, 2022). Our findings suggest decreased SST neuropeptide transmission in PFC circuits may underlie deficits in top-down control of aversive processing. Consistent with this hypothesis, intraventricular infusion of exogenous SST or direct infusion of SSTR2/3 agonists into the mPFC produced antidepressant-like or anxiolytic effects (Brockway et al., 2023; Engin et al., 2008; Engin & Treit, 2009). Taken together, understanding SST neuropeptide transmission in modulating PFC circuits not only enriches the theoretical framework of neuropeptide action within cortical circuits but also opens new avenues for understanding mental health dysfunction and therapeutic interventions.

## Figure Legends

**Figure S1:** Related to figure 1: mPFC^SST^ knockdown impairs cued auditory threat discrimination. (A) Experimental timeline for cued threat discrimination task. (B) Freezing during the baseline session in control and mPFC^SST^-cKO mice (Two-way ANOVA, Genotype x Trial Interaction, *p*=0.8222). (C,D) Freezing during conditioning day 1 *(C;* Two-way ANOVA, Genotype x Trial Interaction, *p*=0.0988) and conditioning day 2 *(D;* Two-way ANOVA, Genotype x Trial Interaction, *p*=0.8403) of cued threat discrimination in control and mPFC^SST^-cKO mice. (E) Test day freezing during the ITI in control and mPFC^SST^-cKO mice (Two-way ANOVA, Genotype x Trial Interaction, *p*=0.7973). (F,G) Similar changes in speed evoked by footshocks during conditioning day 1 (*E;* Two-way ANOVA, Genotype x Trial Interaction, *p*>0.0001) and conditioning day 2 (*G;* Two-way ANOVA, Genotype x Trial Interaction, *p*=0.5047) of threat discrimination in control and SST-loxP mice. (H,I) Open field total distance traveled (*H*; Unpaired t-test *p*=0.5034) and center time (I; Unpaired t-test *p*=0.8764). (J,K) Elevated plus maze time in closed arms (*J*; Unpaired t-test *p*=0.2059) and open arms (*K*; Unpaired t-test *p*=0.8695). (L) Percent time spent in the light zone in the light-dark box assay (Unpaired t-test *p*=0.2266). (M) von Frey filament threshold in grams (g; Unpaired t-test *p*=0.5318). (N) Cold plate withdrawal latency in seconds (s; Unpaired t-test *p*=0.6222). (O) Hotplate lick/jump latency in seconds (s; Unpaired t-test *p*=0.8107). (P) Discrimination index (CS^−^/CS^+^ test day freezing) in SST control (white), mPFC^SST^-cKO (green), vGAT-cKO control (light blue), and vGAT-cKO (blue) during test day (Two-way ANOVA, Manipulation x Transmitter Interaction, *p*=0.4267, Manipulation Main Effect, \*\**p=*0.0027; Tukey’s Post Hoc test, *p*=0.0746 SST-loxP vs vGAT-cKO).

**Figure S2:** related to figure 2 mPFC SST peptidergic regulation of discrimination learning of novel and appetitive outcomes. (A,B) Percentage of active lever (active lever presses / (active + inactive lever presses)) during FR1 (*A*; Two-way ANOVA, Days Main Effect \**p*=0.0036, Genotype Main Effect \**p*=0.0148) and FR5 (Two-way ANOVA with Tukey’s Post Hoc test, \**p*=0.0168 WT lever accuracy vs mPFC^SST^-cKO lever accuracy day 1; \**p*=0.0300 WT lever accuracy vs mPFC^SST^-cKO lever accuracy day 4) sessions in control and mPFC^SST^-cKO mice. (C) Freely available chocolate pellets consumed during exposure to chocolate pellets in the operant chamber without the active and inactive levers (Unpaired t-test *p*=0.4801). D) Total freely available laboratory chow and operant-derived chocolate pellets in grams (g; One-way ANOVA with Tukey’s Post Hoc test, \*\*\*\**p*<0.0001 WT Ad-libitum vs WT FR5, \*\**p*=0.0165 mPFC^SST^ Ad-libitum vs mPFC^SST^ FR5). E) Sucrose preference in WT and SST-loxP mice (Unpaired t-test, \*\*\*\**p*<0.0001).

**Fig. S3:** Related to Figure 3. mPFC^SST^ interneurons encode footshock outcomes, neutral and threat-predictive cues, and post-threat periods. (A,K) CS modulation of mPFC^SST^ neuron activity across CS^+^ and CS^−^ tone presentations during the baseline session (*A*; Two-way ANOVA, Time Main Effect, \**p*=0.0136) and conditioning day 2 (*K*; Two-way ANOVA, Time Main Effect, \*\*\*\**p* <0.0001). (B, O) Z-Scored Ca^2+^ activity in CS^+^ and CS^−^ modulated neurons across all trials during the baseline session (B; Two-way ANOVA, Time x Stimulus Interaction \**p*<0.0212) and conditioning day 2 (O; Two-way ANOVA, Time x Stimulus Interaction \*\*\*\**p*<0.0001). (C,M) Percentage of neurons whose activity is modulated during CS_n_ and also subsequently modulated in proceeding trials CS_n+I_ during baseline (C) and conditioning day 2 (M). Mean percentage of neurons modulated in subsequent trials in baseline day (C; Paired t-test, *p*=0.4529) and conditioning day 2 (M; *p*=0.9015). (D,N) Same as C but for neurons whose activity was modulated during CS_n_ and during preceding trials CS_n-I._ Mean percentage of neurons modulated in subsequent trials in baseline day (D; Paired t-test, *p*=0.1369) and conditioning day 2 (N; *p*=0.3409). (E, Q) Percentage of neurons whose activity was modulated during the ITI following a CS^+^ (ITI^+^) and CS^−^ (ITI^−^) during the baseline session (Two-way ANOVA, time main effect \**p*=0.0484) and conditioning day 2 (Two Way ANOVA, time main effect \*\*\*\**p<*0.0001). (F, R) Ca^2+^ activity in ITI^+^ and ITI^−^ modulated neurons collapsed across trials during baseline session (F; Two-way ANOVA, Time x ITI Type Interaction, *p*=0.8835) and conditioning session day 2 (R; Two-way ANOVA, Time x ITI Type Interaction, \*\*\*\**p*<0.0001). (G,S) Percentage of neurons modulated by speed during the baseline session (G) and conditioning day 2 (S). (H,T) Speed modulated neurons did not respond to changes in speed in a linear manner during the baseline session (H) and conditioning day 2 (T; See statistics table for Pearson Correlation coefficients and *p* values for individual mice). (I) Percentage of mPFC^SST^ neurons modulated by footshocks during conditioning day 2. (J) Z-Scored GCaMP responses aligned to footshocks collapsed across trials and heatmap of Ca^2+^ responses across trials during conditioning day 2. (L) Cumulative probability of CS modulated neurons demonstrating that most CS modulated neurons are recruited in early trials (Two-way ANOVA, Stimulus Main Effect *p=*0.5215s). (P) Z-Scored Ca^2+^ activity in SST interneurons whose activity was modulated during early CS^+^ (trials 1-3) and late CS^+^ during conditioning day 2 (trials 8-10; Two-way ANOVA, Time x Early vs Late Interaction \**p*=0.0298).

**Figure S4:** Related to Figure 4. mPFC SST release during cued auditory threat discrimination promotes discriminative learning. A) GRAB_SST_-mediated fluorescence during presentation of the CS^−^ (left) and CS^+^ (right) during baseline day. AUC of the time windows for the CS^+^ and CS^−^ (top; Paired t-test *p*=0.4004) and the post-CS period (bottom; Paired t-test *p*=0.8853). B) Same as in *A* but during conditioning day 2. AUC of the time windows for the CS^+^ and CS^−^ (top; Paired t-test *p*=0.7987) and post-CS periods (bottom; Paired t-test *p*=0.3343). C) Time course of GRAB_SST_ fluorescence in response to CNO and saline treatment in mice expressing AAV-GRAB-SST and AAV-FLEX-mCherry (Paired t-test *p*=0.8510). D) Footshock-evoked responses in mCherry-expressing control mice and HM3Dq expressing to demonstrate control mice had functional GRAB_SST_ expression. AUC of the time windows for the CS^+^ and CS^−^ (top; Paired t-test *p*=0.8422) and post-CS periods (bottom; Paired t-test *p*=0.8146). E-H) Freezing during CS^+^ and CS^−^ during threat discrimination conditioning day 1 (*E;* Two-way ANOVA, Genotype x Trial Interaction, *p*=0.2138) and conditioning day 2 (*F;* Two-way ANOVA, Genotype x Trial Interaction, *p*=0.0882) in mice injected with Cyclo-SST during conditioning day 1. Freezing during CS^+^ and CS^−^ during threat discrimination conditioning day 1 (*G;* Two-way ANOVA, Genotype x Trial Interaction, *p*=0.8125) and conditioning day 2 (*H;* Two-way ANOVA, Genotype x Trial Interaction, *p*=0.5053) in mice injected with Cyclo-SST during test day. I) Test day freezing during the ITI in mice treated with Cyclo-SST on conditioning day 1 (Two-way ANOVA, Genotype x Trial Interaction, \**p*=0.02237). J) Test day freezing during the ITI in mice treated with Cyclo-SST on test recall day (Two-way ANOVA, Genotype x Trial Interaction, *p*=0.5360).

**Figure S5:** Related figure 5: mPFC SST knockdown alters mPFC neuronal encoding in a cued-threat discrimination task. (A,C,M,N) Percentage of total neurons modulated by CS^+^ (A), CS^−^ (C), ITI (M), and speed (N) during baseline day, WT vs SST-loxP (A, Two-way ANOVA, Time x Genotype Interaction *p*=0.1422; C, Two-way ANOVA, Time x Genotype interaction *p*=0.6835; M, Two-way ANOVA Time x Genotype Interaction *p*=0.8999; N, Unpaired t-test, *p*=0.1694). (B,D) Timecourse of Z-scored GCaMP7f activity of neurons modulated by CS^+^ (B), and CS^−^ (D), during baseline day in WT and SST-loxP. Heatmaps representing Z-scored activity in CS^+^ and CS^−^ across trials in WT (top) and SST-loxP (bottom) mice. (B; Two-way ANOVA, time x genotype interaction *p*= 0.1486, D; Two-way ANOVA, Time x Genotype Interaction *p*=0.0691). (E,F,I,J) Heatmap representing the percent of neurons modulated by CS^+^ (A) or CS^−^ (C) tones across baseline day in WT (left) and SST-loxP (right) mice. Re-modulation percentage was calculated by determining the percentage of neurons with significant modulation in a specific trial (CS^+^_n_ or CS^−^_n_) that were also significantly modulated in n-proceeding (CS_n+i_) trials. % Re-modulation = (CS_n_ | CS_n±i_ / CS_n_), (F, Unpaired t-test *p*=0.3808; J, Unpaired t-test *p*=0.1353). (G,H, K, L) Heatmap representing the percent of neurons modulated by CS^+^ (I) or CS^−^ tones across baseline day in WT (left) and SST-loxP (right) mice. Re-modulation percentage was calculated by determining the percentage of neurons with significant modulation in a specific trial (CS^+^_n_ or CS^−^_n_) that were also significantly modulated n-preceding (CS_n-i_) trial (CS_n_ | CS_n±i_) trials. % Re-modulation = (CS_n_ | CS_n±i_ / CS_n_), (H, Unpaired t-test *p*=0.5974; L, Unpaired t-test *p*=0.2882).

**Figure S6:** Related figure 5: mPFC SST knockdown alters mPFC neuronal encoding in a cued-threat discrimination task. (A,C,O,Q,R) Percentage of total neurons modulated by CS^+^ (A), CS^−^ (C), footshock (O), ITI (Q), and speed (R) during conditioning day 2 in WT and SST-loxP mice (A, Two-way ANOVA, Time x Genotype Interaction \*\*\*\**p*<0.0001; C, Two-way ANOVA, Time x Genotype Interaction \*\*\**p*=0.0003; O, Unpaired t-test, \*\**p*=0.0031; Q, Two-way ANOVA Time x Genotype Interaction \*\*\*\**p*<0.0001; R, Unpaired t-test, *p*=0.6603). (B,D,P) Timecourse of Z-scored GCaMP7f activity of neurons modulated by CS^+^ (B), and CS^−^ (D), and footshock (P) during conditioning day 2 in WT and SST-loxP mice. Heatmaps representing Z-scored activity in response to CS^+^ and CS^−^ across trials in WT (top) and SST-loxP (bottom) mice. (B, Two-way ANOVA, Time x Genotype Interaction \*\*\*\**p*<0.0001; D, Two-way ANOVA, Time x Genotype Interaction \*\*\**p*=0.0008; P, Two-way ANOVA, Time x Genotype Interaction \*\*\**p*=0.0003). (E, H) Cumulative probability of cells that become significantly modulated by the CS^+^ (E) and CS^−^ (H) across discrimination day 2 (E; Two-way ANOVA, Genotype Main Effect \*\*\**p*=0.001; H, Two-way ANOVA, Genotype Main Effect \*\**p*=0.0042). (F,G, I, H) Heatmap representing the percent of neurons modulated by CS^+^(F,G) or CS^−^ (I,J) tones across conditioning day 1 session in WT (left) and SST-loxP (right) neurons. Re-modulation percentage was calculated by determining the percentage of neurons with significant modulation in a specific trial (CS^+^_n_ or CS^−^_n_) that were also significantly modulated in n-proceeding (CS_n+i_) or n-preceding (CS_n-i_) trials (CS_n_ | CS_n±i_). % Re-modulation = (CS_n_ | CS_n±i_ / CS_n_) (K, L) Percentage of neurons with significant modulation in *n*-proceeding trials WT vs SST-loxP (K, Unpaired t-test \*\**p*=0.0065; L, Unpaired t-test \**p*=0.0191). (M, N) Percentage of neurons with significant modulation in *n*-preceding trials WT vs SST-loxP (M, Unpaired t-test *p*=0.2296; N, Unpaired t-test *p*=0.7899).

**Figure S7:** Related figure 5: mPFC SST knockdown alters mPFC neuronal encoding in a cued-threat discrimination task. (A, C, M, N) Percentage of total neurons modulated by CS^+^ (A), CS^−^ (C), ITI (M), and speed (N) during test day in WT and SST-loxP mice (A, Two-way ANOVA, Tone x Genotype Interaction \*\**p*=0.0050; C, Two-way ANOVA, Tone x Genotype Interaction \**p*=0.0278; M, Two-way ANOVA Tone x Genotype Interaction \*\*\**p*=0.0003; N, Unpaired t-test, *p*=0.3019). (B,D) Timecourse of Z-scored GCaMP7f activity of neurons modulated by CS^+^ (B) and CS^−^ (D) during test day in WT and SST-loxP mice. Heatmaps representing Z-scored activity in response to CS^+^ and CS^−^ across trials in WT (top) and SST-loxP (bottom) mice. (B; Two-way ANOVA, Time x Genotype Interaction \*\*\*\**p*<0.0001; D, Two-way ANOVA, Time x Genotype Interaction *p*=0.2210). (E,F,I,J) Heatmap representing the percent of neurons modulated by CS^+^ (E) or CS^−^ (I) tones across test day in WT (left) and SST-loxP (right) neurons. Re-modulation percentage was calculated by determining the percentage of neurons with significant modulation in a specific trial (CS^+^_n_ or CS^−^_n_) that were also significantly modulated in n-proceeding (CS_n+i_) trials. % Re-modulation = (CS_n_ | CS_n±i_ / CS_n_), (F, Unpaired t-test *p*=0.0975; J, Unpaired t-test \**p*=0.0436). (G,H, K, L) Heatmap representing the percent of neurons modulated by CS^+^ (G) or CS^−^ (K) tones across test day in WT (left) and SST-loxP (right) neurons. Re-modulation percentage was calculated by determining the percentage of neurons with significant modulation in a specific trial (CS^+^_n_ or CS^−^_n_) that were also significantly modulated n-preceding (CS_n-i_) trials (CS_n_ | CS_n±i_). % re-modulation = (CS_n_ | CS_n±i_ / CS_n_), (H, Unpaired t-test *p*=0.2654; L, Unpaired t-test *p*=0.0856).

**Figure S8:** Related to Figure 6: mPFC SST knockdown alters encoding of task variables in a threat discrimination task. (A,B) Baseline day data representing the percentage of tones encoding the CS^+^, CS^−^, or both tones across blocks of CS^+^/CS^−^ pairs in WT mice (A, Two-way ANOVA, Encoding Type Main Effect, \*\*\**p=*0.0002) and SST-loxP mice (B, Two-way ANOVA, Encoding Type Main Effect, \*\*\*\**p<*0.0001). (C) Average CS^+^, CS^−^, and encoding of both CS collapsed across presentations in WT and SST-loxP mice (Two-way ANOVA, Encoding Type Main Effect, \*\*\*\**p<*0.0001; Tukey’s Main Effects Comparison; CS^+^ vs both, \*\*\*\**p<*0.0001, CS^−^ vs both, \*\*\**p=*0.0002). (D) Heatmap representing the percent of neurons activated during both the CS^+^ and CS^−^ tones across the baseline day session in WT (top) and SST-loxP (bottom) neurons. Co-modulation percentage is calculated as the number of neurons with significant modulation to CS^+^ in a given trial and all CS^−^ trials. (E) UpSet visualization of neurons modulated by different combinations of footshock, CS^−^, CS^+^ or ITI following a CS^+^ (ITI^+^) and CS^−^ (ITI^−^). Categories with <1% of neurons modulated are not shown. Left horizontal bars represent percentages of cardinal categories, including those not shown. Dashed lines indicate separation of categories encoding different numbers of variables. (F) Heatmaps representing Pearson’s correlation of neural activity across the baseline day session for WT and SST-loxP mice. Red values indicate higher correlation, and blue values indicate lower correlation. (G) Pearson’s correlation of activity from CS^+^ tones 1-3 to the last tone, as represented in F (Two-way ANOVA, Genotype Presentation Main Effect *p*=0.7654). (H) Pearson’s correlation of activity from CS^−^ tones 1-3 to the last tone, as represented in F (Two-way ANOVA, Genotype Presentation Main Effect *p=*0.3668). (I,J) Test day data representing the percentage of tones encoding the CS^+^, CS^−^, or both tones across blocks of CS^+^/CS^−^ pairs in WT mice (I, Two-way ANOVA, Encoding Type Main Effect, *p=*0.2015) and SST-loxP mice (J, Two-way ANOVA, Encoding Type Main Effect, \*\*\*\**p<*0.0001). (K) Average individual and joint CS encoding in WT and SST-loxP mice during test day (Two-way ANOVA, Encoding Type x Genotype Interaction, \**p*=0.0259; Tukey’s Post-hoc test, WT CS^+^ vs CS^−^ \*\**p=*0.0047, SST-loxP CS^+^ vs both, \*\**p=*0.0056, SST-loxP CS^−^ vs both, \*\**p=*0.0048). (L) Heatmap representing the percent of neurons activated during both the CS^+^ and CS^−^ tones across the discrimination day 2 session in WT (top) and SST-loxP (bottom) neurons. Co-modulation percentage is calculated as the number of neurons with significant modulation to CS^+^ in a given trial and all CS^−^ trials. (M) UpSet visualization of neurons modulated by different combinations of footshock, CS^−^, CS^+^ or ITI following a CS^+^ (ITI^+^) and CS^−^ (ITI^−^). Categories with <1% of neurons modulated are not shown. (N) Heatmaps representing Pearson’s correlation of neural activity across the test day session for WT and SST-loxP mice. Red values indicate higher correlation and blue values indicate lower correlation. (O) Pearson’s correlation of activity from CS^+^ tones 1-3 to the last tone (Two-way ANOVA, Genotype Main Effect *p*=0.2559). (P) Pearson’s correlation of activity from CS^−^ tones 1-3 to the last tone (Two-way ANOVA, Genotype Main Effect *p*=0.1562).

**Figure S9:** Related to Figure 6: mPFC SST knockdown alters encoding of task variables in a threat discrimination task. (A,B) Conditioning day 2 data representing the percentage of tones encoding the CS^+^, CS^−^, or both tones across blocks of CS^+^/CS^−^ pairs in WT mice (A, Two-way ANOVA, Encoding Type Main Effect, *p*=0.1252) and SST-loxP mice (B, Two-way ANOVA, Encoding Type Main Effect, \*\*\**p=*0.0005). (C) Average CS^+^, CS^−^, and encoding of both CS collapsed across presentations in WT and SST-loxP mice (Two-way ANOVA, Encoding Type Main Effect, \**p=*0.0220). (D) Percentage of neurons modulated by the CS^+^ and CS^−^ tone across the conditioning day 2 session in WT (left) and SST-loxP (right) mice. Co-modulation percentage is calculated as the number of neurons with significant modulation to CS^+^ in a given trial and all CS^−^ trials. (E) UpSet visualization of neurons modulated by different combinations of footshock, CS^−^, CS^+^ or ITI following a CS^+^ (ITI^+^) and CS^−^ (ITI^−^). Categories with <1% of neurons modulated are not shown. Left horizontal bars represent percentages of cardinal categories, including those not shown. Dashed lines indicate separation of categories encoding different numbers of variables. (F, G) Percentage of neurons with configural encoding (≥4 variables; F, left; Two-sided Fishers Exact Test \*\*\**p*=0.0005) or with limited encoding (≤3 variables; G, right; Two-sided Fishers Exact Test *p*=0.0603) including and excluding footshock in WT and SST-loxP mice. (H) The variable fold difference between WT and SST-loxP mice in distinct categories (each represented by a dot) containing (red) or lacking a footshock (gray) with configural encoding (≥4 variables; left) or limited encoding (≤3 variables; right) of variables. Fold difference is the percentage of neurons encoding a distinct category in WT mice divided by the percentage of neurons in SST-loxP mice. Fold differences greater and less than 1 indicate more encoding in WT and SST-loxP mice for a category, respectively (Two-way ANOVA, Variables x Footshocks Interaction, *p=*0.7812, footshock main effect, \*\*\**p=* 0.0002; Bonferroni Post-hoc analysis, 4-6 variables with footshock vs no footshock \**p=* 0.0369, 1-3 variables with footshock vs no footshock \*\**p=* 0.0025). (I-K) Mean cumulative sum during conditioning day 2 of variables encoded across trial blocks consisting of a CS^+^, ITI^+^, CS^−^, and ITI^−^, in cells with configural encoding (≥4 variables) with (I) or without footshock (J), or in cells with limited encoding (≤3 variables) without footshock (K). Dashed line represents theoretical maximum number of variables that can be encoded (I; Two-way ANOVA, Genotype Main Effect \**p=*0.0120; J; Two-way ANOVA, Genotype Main Effect \**p*=0.0187; K, Two-way ANOVA, Genotype Main Effect \*\**p*=0.0020). (L) Heatmaps representing the mean Pearson’s correlation coefficients of vectorized population Ca^2+^ activity across all time points during conditioning day 2 for WT (left) and SST-loxP mice (right). Black dashed lines indicate shock onset, purple and grey squares on borders indicate CS^+^ and CS^−^ presentations, respectively. (M-R) Mean Pearson’s correlation of activity around CS^+^ (M), CS-(N), footshock (O), ITI (P), CS^+^ to final tone (Q), and CS^−^ to final tone (R) (M, Two-way ANOVA, Genotype Main Effect \*\**p=*0.0089; N, Two-way ANOVA, Genotype Main Effect **0.0119; O, Two-way ANOVA, Genotype Main Effect *p*=0.2642; P, Two-way ANOVA, Genotype Main Effect \*\**p=*0.0054; Q, Two-way ANOVA, Genotype Main Effect \**p=*0.0236; R, Two-way ANOVA, Genotype Main Effect **p=0.0092). (S) Representative principal component analysis (PCA) trajectories of WT (top) and SST-loxP (bottom) throughout the entirety of conditioning day 1. Distinct task epochs are indicated by color. Early to late trials are indicated by light to dark color gradients, respectively. CS^+^ presentations in purple, CS^−^ in blue, and footshocks in red, with baseline period in solid black and ITI in solid grey. Centroid distances (top right) of WT (left) and SST-loxP (right), blue square indicates baseline centroid, red square indicates post-footshock centroid, respectively). (T) Mahalanobis Distance of WT and SST-loxP mice (Unpaired t-test, \**p=*0.0425).

**Figure S10:** Related to Figure 6: mPFC SST knockdown alters encoding of task variables in a threat discrimination task. (A-D) Mean CS^+^/ CS^−^ co-modulation values across all trials for encoding cells represented in co-modulation heatmap for proceeding and preceding trials during baseline (A; Fig. S8D; Two-way ANOVA, Proceeding vs Preceding x Genotype Interaction, *p=*0.8868), conditioning day 1 (B; Fig. 6D; Two-way ANOVA, Proceeding vs Preceding x Genotype Interaction, \**p=*0.0180; Tukey’s Post-hoc test WT vs SST-loxP Proceeding, \*\*\**p*=0.0005, WT Proceeding vs Preceding, \*\*\**p*=0.0086), conditioning day 2 (C; Fig. S9D; Two-way ANOVA, Proceeding vs Preceding x Genotype Interaction, *p=*0.1545) and test day (D; Fig. S8L; Two-way ANOVA, Proceeding vs Preceding x Genotype Interaction, *p=*0.2758; Proceeding vs preceding main effect \**p*=0.0136, Tukey’s Post-hoc test SST-loxP Proceeding vs Preceding, \*\**p*=0.0092) in WT and SST-loxP mice. (E-H) Pearson’s correlation of activity from CS^+^ tones to the first tone during baseline (E, Two-way ANOVA, Genotype Main Effect *p*=0.2047), discrimination day 1 (F, Two-way ANOVA, Genotype Main Effect *p*=0.2015), discrimination day 2 (G, Two-way ANOVA, Genotype Main Effect *p*=0.1029), and test day (H, Two-way ANOVA, Genotype Main Effect *p=*0.8850). (I-L) Pearson’s correlation of activity from CS^−^ tones to the first tone during baseline (I, Two-way ANOVA, Genotype Main Effect *p*=0.3787), discrimination day 1 (J, Two-way ANOVA, Genotype Main Effect *p*=0.3842), discrimination day 2 (K, Two-way ANOVA, Genotype Main Effect *p*=0.1291), and test day (L, Two-way ANOVA, Genotype Main Effect *p=*0.1422).

## Author Contributions

M.A.A., N.R. and H.A.T. designed the study and wrote the manuscript. M.A.A., N.R., R.J.F., H.W., B.B.A. H.A.T. performed *in-vivo* single-cell experiments and analyses. M.A.A. N.R., W. R. M., H.B.R. S.Z., Z.W. and J.E.T performed surgeries and behavioral tests. H.E.Y. R. K. and M.A. validated GRAB_SST_ sensor and performed *in-vivo* recordings. Y.L. generated GRAB_SST_ sensor and provided feedback on study. X.G. performed nociceptive testing. H.A.T supervised the work and acquired resources.

## Acknowledgements

The authors would like to thank members of the Unit on Neuromodulation and Synaptic Integration, Laboratory of Neural Circuits and Behavior, Dr. Christopher Moore, and Dr. Yeka Aponte for their critical feedback on this work. This work was supported by a Brain and Behavior Research Foundation Young Investigator Award (HAT), the NIH BRAIN Initiative (1UF-1NS133763-01), and the NIMH Intramural Research Program. H.E.Y, R.J.F., and H.B.R. were supported by a fellowship from the NIH Center for Compulsive Behavior. R.J.F. was supported by a NIH Postdoctoral Research Associate Training Award. The authors would like to thank Dr. Mark Hoon (NIDCR) for sharing the SST-loxP mice, sharing the vGAT-CRISPR construct, providing constructive feedback on the project, and reviewing the manuscript. The authors would also like to thank Grace Smith for her technical assistance in the study, the NIMH Rodent Behavioral Core, and the NIMH Transgenic Core for their support.

## Disclosures

The authors have no financial conflicts to disclose.

## Notes

### Competing Interest Statement

The authors have declared no competing interest.

