## Extended Methods for "Neuropeptidergic transmission shapes emergent properties of prefrontal cortical circuits underlying learning"

**Experimental subjects**

Adult male and female mice (aged 3-6 months at the start of experiments) were used in this study. There were no significant differences between sexes, data were pooled across sex. All mouse lines were crossed with wild-type C57/BL6J mice. Somatostatin loxP (SST-loxP) transgenic mice were re-derived by the NIMH Transgenic Core (Huang et al., 2018) and used for anatomical characterizations, behavioral assays, and single-cell imaging studies. Heterozygous SST-IRES-Cre mice and homozygous SST-loxP mice were used for fiber photometry and single cell Ca2+ imaging studies. Lastly, Wild-type C57/BL6J mice (WT; C57BL/6J, Jackson Laboratories) were used in all experiments. Mice were group housed in controlled facilities where temperature was set at (70-74 ºF) and humidity set at (40-65%). Mice were maintained on a reverse 12-h light/12-h dark cycle with lights off at 9 am. Where explicitly noted, mice were dual-housed with a perforated Plexiglas divider down the middle of the home cage. This was necessary for experiments that required food restriction, and/or protection of cannula, miniaturized endoscopes, and/or fiber optic implants. Mice had *ad-libitum* access to laboratory chow except for cases where animals underwent food restriction procedures. Water was available *ad-libitum*. All procedures were performed per the National Institutes of Health Guide for the Care and Use of Laboratory Animals and approved by the Animal Care and Use Committee of the National Institute of Mental Health Intramural Research Program. All efforts were made to minimize pain, distress, and the number of mice used.

**Stereotaxic surgeries and optical fiber/guide cannula implantation**

Surgeries were conducted under aseptic conditions and body temperature was maintained at approximately ~36 ºC with an electronic heating pad. All mice undergoing stereotaxic surgeries, 8–16 weeks of age, were anesthetized with a mixture of Ketamine (100 mg/kg; ip) and Xylazine (10 mg/kg; ip). The absence of a flinching response to a light pinch indicated an animal’s anesthetic status. The animal’s head was shaved using an electric razor, and ophthalmic ointment (GenTeal) was applied to the eyes to prevent drying and irritation. Mice were then placed in a stereotaxic apparatus (David Kopf Instruments Model 1900, Tujunga, CA, USA). The animal's head was subsequently cleaned with povidone-iodine and 70% ethanol repeatedly. The surgical site was exposed using a sterile scalpel. The mouse’s head was leveled by ensuring the difference in dorsoventral distance between bregma and lambda was within 30 µm. A small craniotomy (diameter ≈ 300 µm) was made above the injection site with a stereotax-mounted drill. The following injection coordinates (in mm) were used for targeting the mPFC (AP: +1.65; ML: ±.3; DV -2.2), motor cortex (AP: 1.5, ML: ±1, DV: -1), and auditory cortex (AP: -2.8, ML: ±3.9, DV: -2.2). Viral infusions of 200-400 nL / site were made at a rate of 100 nL/min utilizing 33 Ga microinjection needles connected to polyethylene 20 (PE20) tubing attached to a 2 µl Hamilton syringe controlled by a microinjection pump (UMP3, World Precision Instruments). After infusion, injectors were kept in place for 8 min to allow diffusion of viral particles and then were slowly withdrawn. Following all surgical procedures that did not require an optical fiber or GRIN lens implant, incisions were secured using VetBond (3M). After the surgery, mice recovered from anesthesia in electronic heating pads until being transferred back to their home cage. All animals that underwent surgical procedures received subcutaneous injections of Ketoprofen (5 mg/kg body weight) for three consecutive days (post-operative analgesia and anti-inflammatory actions). Experiments involving the use of AAVs were performed 3-6 weeks after injection.

For behavioral and anatomical experiments involving genetic ablation of SST in the mPFC, WT or homozygous SST-loxP mice received bilateral injections of AAV8-hSyn-Cre-GFP (300 nL).

For fiber photometry experiments, WT or SST-Cre mice were unilaterally injected with either 300 nL of AAV2-hSyn-GRAB_SST2.0_, and an optical fiber (Doric Lenses; 400 µm; NA 0.66, B280-4604-4.1) was placed on an optical fiber holder mounted on the stereotax and were inserted 100 µm above the viral infusion area (DV -2.1). Optical fibers were secured to the animals using Metabond cement (Parkell, Inc.) with additional layers of dental cement (Lang Dental Manufacturing). For chemogenetic activation of SST interneurons to activate GRAB_SST2.0_, mice were injected with AAV5-hSyn-hM3Dq-mCherry or AAV2-hSyn-mCherry 2 weeks prior to AAV2-hSyn-GRAB_SST2.0_ infusions. To investigate the impact of dmPFC^SST^ genetic deletion on GRAB_SST2.0_ responses, WT or SST-loxP mice first received bilateral injections of AAV8-hSyn-Cre-2A-tdTomato (300 nL), Two weeks later AAV2-hSyn-GRAB_SST2.0_ (300nL respectively) was infused.

For *in-vivo* Ca^2+^ recordings, WT,SST-loxP, or SST-Cre mice first received bilateral injections of AAV8-hSyn-Cre-2A-tdTomato (300 nL), 2 weeks later AAV9-Syn-jGCaMP7f (300nL) was infused. A gradient refractive index (GRIN) lens (Inscopix; 1 x 4 mm) was inserted 200 µm above the viral injection site (DV -2.1). GRIN lens was secured to the skull using Metabond cement (Parkell, Inc.) with additional layers of acrylic dental cement (Lang Dental Manufacturing).

For cannula/drug microinjection experiments, mice were implanted with stainless steel guide cannulas (26 gauge, P1 Technologies) 1 mm above the mPFC (AP: +1.65; ML: ±.3; DV -1.2 from dura). Guide cannulas were secured to the skull using Metabond cement (Parkell, Inc.) with additional layers of dental cement (Lang Dental Manufacturing). Dummy cannulas (P1 Technologies) whose tips were flush with the guide cannulae were used to prevent clogging or introduction of external debris.

**Histology**

After completion of experiments, mice were anesthetized with Euthanasia solution (pentobarbital sodium and phenytoin sodium active ingredients; VedCo Inc.), and transcardially perfused with 25 mL of phosphate-buffered saline (PBS, pH 7.4), followed by 25 mL of 4% paraformaldehyde (PFA) mixed in PBS. Brains were then extracted and post-fixed in 4% PFA at 4 °C overnight, then transferred to PBS for 24 hours until sectioning on a vibratome (Leica VT1200s). For vibratome preparations, brains were embedded on the mounting disk with lab glue and placed in cold PBS bath under the sectioning portion of the vibratome. Tissue was sectioned into 50- or 100-μm thick slices for viral validation, GRIN lens, and/or optical fiber implant verification. Slices were mounted on slide glasses with DAPI Fluoromount-G mounting medium (Southern Biotech, Cat. 0100-20). Confocal or widefield images from all injected and/or implanted mice were examined to determine whether viral expression was expressed in the region of interest (dmPFC, auditory, or motor cortex). Only data from mice with accurate transgene expression, GRIN lens, optical fiber, or guide cannula placement in area of interest were included in the study. Excluded animals from data analyses were due to (1) no or weak transgene expression; (2) off-target transgene expression outside the region of interest; and/or (3) inaccurate implant placement.

**Behavior**

All behavioral experiments took place during the animal’s dark cycle (between 9 am and 6 pm). Behavioral testing commenced 4-6 weeks after surgical procedures. Animals were age- and sex-matched, and dual-housed (see above) at least 3 days before the start of any testing. Separate cohorts of mice were used for all *in-vivo* experiments. Mice were acclimated to red-lit and sound-controlled testing rooms for at least 30 minutes before the start of behavioral testing.

**Cued auditory threat conditioning task:** Mice were placed in sound-attenuating chambers (Med Associates). Task procedures lasted four days. On all days, mice were transferred from the vivarium to a holding cage adjacent to the behavioral training area for 30 minutes before behavioral testing. *Day 1:* Mice were exposed to context “A” for 15 minutes where they received six interleaved presentations of two distinct auditory tones (three presentations of each tone; 70-dB 4kHZ or 12 kHZ; 60s intertrial intervals). Context A consisted of a hexagonal environment with a smooth tactile surface and a vinegar scent and cleaned with acetic acid. *Day 2 and 3:* Mice were exposed to context “B” for 40 minutes in which they received 10 presentations of two distinct auditory tones (70-dB 4 or 12 kHZ, 10 each, pseudo-randomly presented, with 30-60s intertrial intervals). During these two days of context B training one of the tones co-terminated with a mild footshock (CS+; 0.4 mA). Co-termination of the other tone was without consequence and therefore a neutral outcome (CS-). Context B consisted of a square environment with a shock grid and was cleaned with 70% ethanol. *Day 4:* Mice were reintroduced to context A for 15 minutes where they received six interleaved presentations of the CS+ and CS-. All days included 2 minutes of baseline prior to the presentation of the first tone. Freezing (2 sec threshold) was assessed using ANY-maze software (Stoetling) as a defensive state elicited by threats under present conditions.

**Microinjection procedures:** Intra-dmPFC microinjections of the SST-receptor antagonist cyclo-somatostatin (cyclo-SST) were used to pharmacologically inhibit dmPFC^SST^ signaling. Two weeks post-surgery, mice were habituated to guide cannula manipulation for 3 days prior to behavioral testing. We bilaterally infused 500 nL of vehicle (PBS) or cyclo-SST (dissolved in PBS) either on the first day of cued threat acquisition in Context B or day 4 cued threat recall in Context A (Fig. 4) using 33-gauge injector cannulas connected to a syringe pump (UMP3, World Precision Instruments) with PE20 tubing. Injection cannulae protruded 1 mm beyond the tip of the 26-gauge guide cannula. Micro-infusions were delivered over a 1-minute period. After infusion, injectors were left in place for 2 min to allow for complete drug diffusion. Mice were then placed in their home cage for 10 minutes. Subsequently, mice were placed in testing chambers. Cannula placements were verified by histology after an injection of 300 nL per hemisphere of red fluorescent Retrobeads (Lumafluor).

**Operant conditioning:** Mice were food restricted for three days before the start of operant training to achieve 85% of their free-feeding body weight. Mice were weighed daily and fed standard laboratory chow after operant sessions accordingly to maintain weight at 85% pre-restriction weight. Animals had *ad-lib* access to water throughout the study. Operant testing lasted for 4 weeks and was conducted 7 days per week in sound-attenuated mouse operant chambers equipped with two retractable levers and a pellet dispenser (Med-Associates, Fairfax, VT, PC 5 software). Active lever pressing resulted in delivery of sucrose chocolate pellets (20 mg, Bio-Serv). Inactive lever presses had no consequence. A 5 second “time-out” period followed chocolate pellet delivery where active lever presses did not trigger reward delivery. Designation of the active or inactive lever to the right lever was counterbalanced across animals. Operant chambers were dark (house light off) until the start of the session. At the start of the session, the house light was illuminated and remained on for the duration of the task. Before the start of operant training, animals underwent three shaping sessions wherein chocolate pellets were delivered at a random interval schedule (45 sec mean; ranging between 4–132 sec). During shaping sessions, neither lever had a consequence on reward delivery. Days 1-3 of operant training consisted of reward delivery on a fixed-ratio 1 (FR1) schedule, wherein animals had to press the active lever once to obtain a reward. On Days 4-10 rewards were delivered on an FR5 schedule, wherein animals were required to press the active lever five times to obtain a reward. FR1 and FR5 days consisted of one session per day. Shaping and FR sessions lasted 45 minutes.

**Operant Choice task:** Upon the completion of FR5 schedule training, animals underwent testing on an operant choice task. The day before the choice task day, animals were exposed to either freely available laboratory chow (chow-only session) or chocolate pellets (chocolate-only session) in the side opposite to the levers in the operant chambers for 45 min. Importantly, during chow-only sessions and chocolate-only sessions, both active and inactive levers were retracted. The chow-only session took place the day before the freely-available laboratory chow / FR5 chocolate choice test. Freely-available chocolate / FR5 chocolate choice test took place one day after the chocolate-only session. During choice tests, mice were allowed to either lever-press to obtain the sucrose chocolate pellets on an FR5 schedule, or to consume either freely available laboratory chow or sucrose chocolate pellets located on the opposite side of the operant chamber (*ad-lib* chow vs FR5 sucrose chocolate pellets or *ad-lib* sucrose chocolate pellets vs FR5 sucrose chocolate pellets). Laboratory chow and chocolate pellet choice sessions were conducted on different days. After each choice session, a retrain FR5 session was administered, to ensure stable active lever pressing. The number of lever presses (active and inactive), the quantity of standard chow or pellet consumed (in grams), and the total amount of food consumed (freely available and operant-derived) were recorded.

**Progressive ratio operant tasks:** Mice were exposed to an FR5 schedule session after the last choice test before starting progressive ratio (PR) schedule testing. Initially, animals were placed in a PR3 schedule of reinforcement, for 3 consecutive days. Under the PR3 schedule, linear increments of three lever presses (3, 6, 9, 12, 15, etc.) were needed for the delivery of sucrose chocolate pellets. After three PR3 days, animals underwent three days of PR7 reinforcement schedule. Under PR7, linear increments by seven lever presses (7, 14, 21, 28, 35, etc.) were needed for the delivery of sucrose chocolate pellets. Only the active lever was present during PR sessions. Total lever presses and breakpoint, the last ratio of responding reached at the end of the session, were recorded. PR3 and PR7 sessions ended after 1 hr and 3 hrs, respectively, or until the animals went 5 minutes without an active lever response (breakpoint).

**Elevated Plus Maze (EPM):** The EPM is a cross-shaped arena that consists of two open arms and two closed arms. The apparatus is elevated above ground (62.23 cm). These compartments of the cross-shaped arena consisted in two open arms (33.02 L x 5.08 W cm) and two closed arms (33.02 L x 5.08 W x 15.24 H cm). Mice were allowed to explore the maze for 5 minutes. ANY-maze tracking was used to monitor distance travel and time spent in arms.

**Light /dark box:** This assay consists of a box with two compartments; an open-lit area (43.815 L x 21.9 W cm) and a closed dark compartment (43.815 L x 21.9 W cm). To toggle between the two compartments mice can cross an entry point (4 W x 5 H cm), and were allowed to explore for 10 minutes. Mice were placed in the open-lit area, and latency to dark, time spent in dark and light areas was recorded using ANYmaze.

**Open Field Test:** Mice were placed in an open-field arena (43.8 L x 43.8 W x 39.4 H cm) to assess general locomotion and anxiety-like behavior. The total distance traveled was measured for 30 minutes. The area was divided into “center” (23 L x 23 W cm) and “edge” zones, and the percentage of time spent moving and time in the center of the arena was assessed using ANYmaze.

**Novel Object Recognition Test:** This test was conducted using a (43.8 L x 43.8 W x 39.4 H cm) chamber. During baseline mice freely explored the apparatus for 10 minutes; subsequently, two identical objects were placed in opposing corners of the arena. Mice are then allowed to interact with the two objects for 20 minutes (test 1). One day later, mice were placed back into the arena and allowed to freely explore the apparatus for 10 minutes. Subsequently, on Test 2, two objects were placed in opposing corners (one novel and one from test 1). Interaction time was defined as time spent in an interaction zone engaging with any of the objects using ANY-maze.

**Von Frey:** Mice were placed in plexiglass enclosures on a wire mesh frame and were allowed to acclimate for at least 60 min before experiments. The plantar surfaces of the hind paws were stimulated with calibrated von Frey filaments starting from a filament weight of 0.4 g. Filaments were gently pressed against the hind paw plantar surface until the filament bent slightly and was held in place for 3-5s. For each trial mice were tested with the same filament 3-5 times, and the number of paw withdrawals was recorded. The weight of the next filament was heavier or lighter than the previous, with higher-weight filaments chosen when <50% of responses failed to produce a paw withdrawal and were lower when >50% of trials resulted in a withdrawal. This process was repeated five times to determine the 50% withdrawal threshold.

**Hot plate:** Mice were placed in a plexiglass container on a heated plate set to 52°C (IITC Life Science, PE34), and time taken for the first defensive response (flicking, shaking, or licking the paw) was measured. Additionally, the time to first jump was taken as a proxy for escape-like behaviors. Mice were removed from the hot plate either after the first escape attempt or after 60s if no attempts were made to escape. An average result from 3-4 trials performed on separate days is reported.

***In-vivo* Fiber photometry**

*In-vivo* fiber photometry recordings were acquired throughout all phases of the cued threat discrimination task. Mice were handled and habituated to the fiber patch cord in the animal's home cage for at least two days before the start of the recording sessions. This was done by attaching the optic fiber to the implanted fiber using a ferrule sleeve (Doric, ZR_2.5). For recordings, mice were connected to the fiber photometry system and behavior was conducted as detailed above. For each session, we recorded video (Anymaze) and fiber photometry signals (Synapse), which were synchronized using TTL. GCaMP fluorescence was assessed by a fiber photometry system (Tucker-Davies Technologies) using a sinusoidally modulated 470 nm (531 Hz) LED (DC4104, ThorLabs) for acquiring Ca^2+^-dependent GCaMP activity and an isosbestic, Ca^2+^-independent, autofluorescence signal excited by a 405 nm LED (211 Hz) that allowed us to correct for movement-related and photobleaching artifacts. The isosbestic channel represents the point of GCaMP excitation at which emissions from the Ca^2+^-bound and unbound states have equal intensities. Thus, the control channel represents GCaMP emissions that are independent of fluctuations in Ca^2+^ concentration, but still susceptible to artifacts related to movement, fiber bending, and all other physical alterations. Signals were sampled by the TDT program Synapse at a sampling rate of 1,017 Hz. Both LEDs were fed into a fluorescent minicube (Doric Lenses) connected to a silicon photoreceiver (Model 2151; Newport) or integrated lock-in amplifiers (Doric Lenses). Optical signals were then collected, digitized at 6 kHz, and recorded using the real-time RZ5P processor (Tucker-Davis Technologies). Data were then acquired using the Synapse software, which controls the RZ5P lock-in amplifier. Optical patch cables were connected to the lasers through a rotary adaptor and were extensively photo-bleached before recordings to reduce auto-fluorescence. At the beginning of each session, light power was adjusted to 40-60 μW at the intersection between the fiber tip and the animal and 470 nm and 405 nm outputs were kept consistent for each mouse.

**Analysis of fiber photometry data**

Photometry analyses was performed with custom scripts written in Python together with the pMAT package in MATLAB (Bruno et al., 2021). The time of tone presentations was defined as time = 0, positioned 5 sec before and 45 sec after tone onset. All 10 CS+ and CS- trials were averaged within mice, with time course of z-scored data representing mean±SEM across animals. AUC for the CS+ or CS- time window were calculated from time frames 0-28 sec from tone onset. The AUC for the footshock response was calculated from 28-45 sec after tone onset. Videos were analyzed with Anymaze.

**Single Cell Ca^2+^ Imaging**

We obtained single-cell Ca^2+^ recordings through an integrated GRIN lens with a head-mounted miniaturized microscope (miniscope; nVoke, Inscopix) approximately 4 weeks after implantation. Each mouse was habituated for three days, in which the miniscope was attached to the baseplate and mice were put in an open field for 10 min. During the habituation, miniscope settings (LED power, gain, and focus position) for each mouse were determined, and these settings were kept fixed for each mouse throughout cued threat discrimination recordings.

To identify individual regions of interest (ROIs) Ca^2+^ dynamics from raw miniscope recordings, imaging data were processed using the Inscopix Data Processing Software (IDPS, Inscopix). Videos were spatially downsampled by a factor of 4 and temporally downsampled to 10 Hz, bandpass filtered (0.005-0.5 pixels), motion corrected, and extracted using constrained nonnegative matrix factorization-extended (CNMFe) using the following parameters: spatial downsample factor = 1, temporal down sample = 1, Gaussian filter size = 2, maximum neuron diameter = 5, minCorr=.8, min pnr=10, merge threshold = 0.8, ring size factor = 1.4, patch size = 80, patch overlap = 20). After CNMFe, neurons were visually inspected by experimenters for clear somatic morphology of appropriate size, shape, and appropriate Ca^2+^ dynamics (faster rise and slower decay; signal to noise ratio). Cells with abnormal morphology, sub-standard Ca^2+^ traces, and/or inadequate spatial separation from other neurons were rejected from the final cell set. The data were then exported to CSV files and custom-written Python scripts were used for analysis.

**Multiple Linear Regression (MLR) with Temporal Kernel**

To determine how the activity of single cells in the mPFC was influenced during cued threat discrimination, along with changes in locomotor activity, we employed a multiple linear regression model. Each neuron was fitted to the following MLR model:

${Ca}^{2+}{Signal}_{t}= \beta_{0}\left( {Ca}^{2+}{Signal}_{t-1} \right) + \sum_{n=1}^{10} \sum_{i=0}^{4} \beta_{1ni}\left( {CS}_{n\left( t-i \right)}^{+} \right)+\sum_{n=1}^{10} \sum_{i=0}^{4} \beta_{2ni}\left( {CS}_{n\left( t-i \right)}^{-} \right)+ \sum_{n=1}^{9} \sum_{i=0}^{4} \beta_{3ni}\left( {ITI}_{n\left( t-i \right)}^{+} \right)+ \sum_{n=1}^{9} \sum_{i=0}^{4} \beta_{4ni}\left( {ITI}_{n\left( t-i \right)}^{-} \right)+ \sum_{i=0}^{4} \beta_{5i}\left( {Footshock}_{t-1} \right)+\sum_{i=0}^{4} \beta_{6i}\left( {Speed}_{t-1} \right)+ \varepsilon$

The dependent variable in this model was the ΔF/F_noise_ value (*Ca*^2+^ *Signal_t_*) of each neuron, and the independent variables ($\beta_{1-5})$were speed and task-related variables, including the CS+, CS-, footshock, intertrial interval (ITI) following CS+ (ITI+) and ITI following CS- (ITI-), with a time kernel. Data were aggregated into 500 ms bins, and the time kernel consisted of 5 time bins (${}_{i}$), equivalent to 2.5 seconds. This allowed us to investigate how past stimuli or events influenced current neural activity, which is crucial for understanding the encoding of information by individual neurons. β_0_ (*Ca*^2+^ *Signal_t_*_−1_) is an autoregressive term to account for the influence of past Ca^2+^ signals on current events. This term included the Ca^2+^ signal lagged by one-time bin (500 ms). Each $\beta$ is the coefficient of this model capturing the strength and direction of the relationship between each independent variable and Ca^2+^ activity in each individual neuron. Trial sequence was accounted for by coding each trial stimulus / ITI as an independent variable: CS^+^_n_, CS^-^_n_, Footshock_n_, ITI^+^_n_, ITI^-^_n_ are stimuli / ITI on trial_n_. To analyze the encoding of stimuli/task phases (CS^+^, CS^-^, ITI^+^, and ITI^-^) on a trial-by-trial basis, a design matrix was constructed with dummy-coded variables to represent the presence or absence of each stimulus. Each stimulus was allocated as a separate column, with time points as rows. Specifically, when the CS^+^, CS^-^, footshock, ITI^+^, or ITI^-^ was present during a particular time point, coded as 1; conversely, when these stimuli were absent during that time point, it was coded as 0. The speed of mice was treated as a continuous variable. Speed and footshocks were not subdivided into individual trials.

We employed diagnostic analyses to assess potential multicollinearity issues in our regression model, including assessing the Variance Inflation Factor (VIF), where values exceeding 10 suggest the presence of multicollinearity, and the Condition Index, where values greater than 30 indicate multicollinearity. A minimum eigenvalue plot that incorporates the newly added predictors to gain insights into how the multicollinearity changes as more predictors are introduced was examined. If the minimum eigenvalues start to drop significantly, indicating increasing multicollinearity, reconsideration of the choice of predictors is necessary. Based on the diagnostic analyses we chose speed instead of freezing in our predictor because freezing was an independent variable that caused high multicollinearity in the model.

To assess the significance of variables in influencing neural activity, we compared the variance explained by the full model, which included the variable of interest, to that of a reduced model that excluded a specific variable, allowing us to test whether the difference in explained variance was statistically significant (p<0.05). A variable that significantly modulated neural activity was considered "encoded" in the neural signal of a given neuron (Costa et al., 2016; Wang et al., 2024). UpSet plots to visualize variable set intersections and cardinality was implemented in Python using the UpSetPlot function (Lex et al., 2014).

The Kullback-Leibler Divergence test was calculated to compare the distribution of proportion of neurons encoding multiple variables in WT and SST-loxP mice using the following:

$$D_{KL}(P \left\| Q \right.) = \sum_{x\in X} P\left( x \right)\log\left( \frac{P(x)}{Q(x)} \right)$$

where $P(x)$ and $Q(x)$ represents the distribution probabilities of neurons with different combinations of variables encoded in the test and reference group, respectively. Since the KL-divergence test result is not symmetrical, the KL divergence was calculated with WT or SST-loxP mice as $Q.$

**Population-level analyses of single-cell Ca^2+^ imaging**

We performed population-level analyses to examine how SST may regulate network dynamics as previously described (Wang et al., 2024). For Ca^2+^ transients from the experiment comparing dmPFC^SST^ cKO versus WT controls, the whole session recording of each cell was standardized (Z-Scored) relative to the first 2 minutes (baseline period) of the recording. We analyzed dynamic changes at the population level (ΔF/F_noise_) using Principal Component Analysis (PCA) to generate the trajectory of the network through the neural state space for each mouse. First, we derived the ΔF/F_noise_ of all neurons at each sampled time point. The matrix dimensions consisted of time points by the number of neurons. The first and second components were plotted for each time point and color-coded according to epoch during the session (i.e., baseline, CS+, CS-, shock, and ITI). To quantify the change in neural activity induced during our cued threat discrimination task, we first labeled each mouse’s PC values at each timestep as being either “baseline” or “post-baseline”. The baseline was defined as the set of all time points up to, but not including, the first shock. Post-baseline was defined as the set of all time points from the onset of the first shock until the end of the task. We then identified the centroids of derived PC coordinates for baseline and post-baseline periods and computed the Mahalanobis distance between them as follows:

$$d_{M}\left( \vec{x}, \vec{y,} Q \right)= \sqrt{\left( \vec{x}- \vec{y} \right)^{T}S^{-1}\left( \vec{x}- \vec{y} \right)}$$

where $\vec{x}$ and $\vec{y}$ represent the centroids at baseline and post-baseline, respectively; *Q* represents the probability distribution of $\vec{x}$ and $\vec{y}$ in $R^{N}$and $S^{-1}$designates the inverse covariance matrix.

To monitor correlated activity across different behavioral epochs we generated a correlation matrix where each n-dimensional (e.g. number of neurons) population vector at time *t* was correlated (mean Pearson’s correlation coefficient (PCC)) with the population vector of the same neurons at every other time point within the session, yielding a t x t matrix which represents the “moment by moment” changes in correlated activity across the population throughout the entire session. We then calculated the mean PCC within or across specific trial epochs of defined windows (28s of the cue, 5 sec during the shock, and the ITI). Because the PCC matrix is symmetric and contains values of 1 along the diagonal, we factored out the diagonal from our computations of the mean PCC at each window.
